# Deciphering Cell-Type and Temporal-Specific Matrisome Expression Signatures in Human Cortical Development and Neurodevelopmental Disorders via scRNA-Seq Meta-Analysis

**DOI:** 10.1101/2025.02.24.639826

**Authors:** Do Hyeon Gim, Muhammad Z.K. Assir, Paul A. Fowler, Michael D. Morgan, Daniel A. Berg, Eunchai Kang

## Abstract

Human cortical development is a complex process involving the proliferation, differentiation, and migration of progenitor cells, all coordinated within a dynamic extracellular matrix (ECM). ECM plays a crucial role in guiding these processes, yet its specific contributions and the implications of its dysregulation in neurodevelopmental disorders (NDDs) remain underexplored. In this study, we conducted a meta-analysis of single-cell RNA sequencing (scRNA-seq) data from 37 donors, gestational weeks (GWs) 8 to 26 across six independent studies to elucidate cell type-specific matrisome gene expression signatures and their dynamics in the developing human cortex. Our analysis identified distinct matrisome gene signatures across various cell types, with significant temporal changes during cortical development. Notably, a substantial proportion of matrisome genes are associated with NDDs, exhibiting cell type, temporal and disease specificity. These findings highlight the critical role of cell type-specific matrisome regulation in cortical development and its potential involvement in NDD pathogenesis. This study provides a comprehensive map of cell type-specific matrisome signatures in the developing human cortex and highlights the importance of ECM in both normal development and the pathogenesis of NDDs.

## INTRODUCTION

Human cortical development involves a series of intricate processes, including the proliferation of progenitor cells, their differentiation into distinct cell types, and their subsequent migration and interconnection. The precise coordination of these processes is crucial for shaping the cortex’s developmental trajectory and defining its final architecture. While considerable progress has been made in understanding the regulation of individual processes, how they are coordinated through simultaneous interactions among different cell types remains poorly understood.

The extracellular matrix (ECM) is of particular interest because it provides a dynamic environment for interactions among various cell types. The ECM serves as the structural framework in which the cellular components of all tissues are embedded. In the developing human brain, the ECM constitutes approximately 40% of the total volume^1^, offering crucial support and guidance for cellular organization and function, and regulating cell-cell communications by governing signaling pathways through controlling spread and restriction of signaling molecules^2,3^. During neural development, the ECM is essential for the proliferation and differentiation of neuronal progenitors^4–7^, dendritic and axonal growth^8–11^ and axonal guidance^11^, migration^8^, cortical folding^12–14^, connectivity, and synaptic plasticity^15–17^.

The importance of the ECM in cortical development is highlighted by neurodevelopmental disorders (NDDs) linked to ECM gene mutations, such as *RELN*, which is essential for neuronal migration and positioning^18,19^. Mutations in ECM-related genes, including *POMT1/2* and laminin subunits, are associated with conditions such as cobblestone lissencephaly and polymicrogyria, leading to cortical layer disorganization and abnormal cortical folding^20–22^. Furthermore, mutations in ECM components like collagen are linked to malformations such as porencephaly and Knobloch syndrome, where neuronal migration is disrupted^23^.

While functional studies of ECM genes in animal models, particularly mice, have deepened our understanding of ECM roles during cortical development, significant differences exist between the ECM of the human and mouse cortex. The human fetal cortex ECM is more abundant and diverse, especially rich in components such as hyaluronan, chondroitin sulfate proteoglycans, and other glycosaminoglycans, whereas the mouse ECM is less complex^24^. These differences are also evident in their transcriptome profiles and gene expression patterns. In the developing human cortex, the ECM transcriptome of the subventricular zone (SVZ) closely resembles that of the ventricular zone (VZ), suggesting a shared microenvironment that supports progenitor cell self-renewal, and includes distinct sets of collagens, laminins, proteoglycans, integrins, and specific growth factors^25^. In contrast, in the developing mouse cortex, the SVZ ECM transcriptome more closely resembles that of the cortical plate (CP), indicating a fundamental difference in the organization and function of germinal zones across species^25^. Gene expression analysis further reveals that ECM-associated genes are highly expressed in both the VZ and SVZ in humans, suggesting a shared ECM environment that supports the self-renewal of neural stem cells and progenitors. In contrast, in mice, these genes are predominantly expressed in the VZ^25^. These differences might contribute to the greater plasticity and complexity of the human cortex.

The matrisome refers to the set of genes and proteins that compose and regulate the ECM^18^. To better understand the developmental processes and their dysregulation contributing to NDD pathogenesis, it is important to systematically examine how the matrisome shapes human cortical development. It is particularly critical to understand how each cell type specifically contributes to the matrisome and mediates diverse biological processes throughout development.

Single-cell RNA sequencing (scRNA-seq) of human fetal brain tissue is a powerful tool that advances our understanding of cell-type-specific gene expression dynamics at single-cell resolution. This technology facilitates the analysis of cellular interactions across various developmental stages, providing detailed insights into the progression of cellular differentiation and development. By capturing snapshots of individual cells at different stages, scRNA-seq offers a comprehensive view of the cellular landscape, revealing the intricate processes underlying brain formation and maturation^26^. This method enhances our ability to study the complex interactions mediated by the matrisome and transitions that occur during neurodevelopment, contributing to a deeper understanding of brain function and the pathogenesis of NDDs.

However, the limited availability of human fetal tissue, due to ethical, legal, and logistical constraints, presents a significant challenge to using scRNA-seq to study dynamic brain development^26^. Consequently, individual studies often rely on a small number of available samples. These limitations prevent findings from fully capturing the diversity and complexity of cellular states and interactions present at each stage of dynamic fetal brain development, leading to gaps in our understanding of a comprehensive and continuous map of brain development.

Meta-analysis of scRNA-seq data from human fetal brain tissue offers substantial benefits in addressing the scarcity of tissue samples across different developmental stages. By integrating datasets from multiple studies, meta-analysis can significantly increase the sample size and reduce biases inherent in individual datasets, providing a more comprehensive overview of gene expression patterns and cellular dynamics throughout brain development^27^. Importantly, meta-analysis facilitates cross-validation of findings, enhancing the reliability and robustness of conclusions drawn from scRNA-seq data^27^.

To elucidate cell type-specific matrisome gene expression signatures and their dynamics across different developmental trajectories in the developing human cortex, we conducted a meta-analysis of scRNA-seq data encompassing gestation weeks (GW) 8 to 26 from six independent studies^28–33^. Our findings reveal that each cell type possesses unique matrisome gene expression signatures, which reflect the biological processes active during cortical development. These signatures undergo dynamic changes along specific differentiation lineages and throughout brain development. Additionally, we discovered that a substantial portion of matrisome genes is associated with NDDs exhibiting cell-type, temporal, and disease-specificity.

## RESULTS

### Matrisome genes linked to NDDs through cross-referenced database analysis

The human matrisome consists of core matrisome proteins, including glycoproteins, proteoglycans, and collagens, as well as matrisome-associated proteins, which are classified into ECM-affiliated proteins, ECM regulators, and secreted factors that bind to the ECM **(Fig. 1a)**. To date, 1,027 proteins have been identified in the human matrisome, comprising 274 core matrisome proteins and 753 matrisome-associated proteins **(Fig. 1a)**^34^. To investigate the association between ECM genes and NDD risk, we cross-referenced ECM genes with three NDD risk gene databases: the Simons Foundation Autism Research Initiative (SFARI) database, the Geisinger Developmental Brain Disorder Gene Database, and the Systems Biology of Neurodevelopmental Disorders (SysNDD) database^35–37^. These databases collectively identified 2,723 unique NDD risk genes, of which 139 are matrisome genes. We found that 17.2% of core matrisome genes and 9.8% of matrisome-associated genes are reported as NDD risk genes **(Fig. 1b)**. Matrisome genes were identified as risk factors for various NDDs, including intellectual disability (ID), autism spectrum disorder (ASD), epilepsy (EP), attention deficit hyperactivity disorder (ADHD), schizophrenia (SCZ), and cerebral palsy (CP) **(Fig. 1c)**. While some core matrisome NDD risk genes, such as *LAMA1, LAMA2*, *RELN*, *COL4A1*, *EYS*, *FBN2*, and *LAMB2*, were linked to multiple NDDs, the majority were associated with a single disorder **(Fig. 1d)**. Similarly, matrisome-associated genes such as *F2*, *FGF13*, *FLG*, *NGLY1*, *SEMA5A*, *CRLF1*, and *FGF14*, were found to be risk factors for more than one type of NDD **(Fig. 1d)**. This finding indicates that both unique and shared matrisome genes are linked to NDDs, suggesting a potential role for their dysregulation in NDD development. Based on these results, we sought to uncover the cell-type-specific expression patterns of matrisome genes and their dynamic changes throughout human cortical development.

**Fig. 1:**
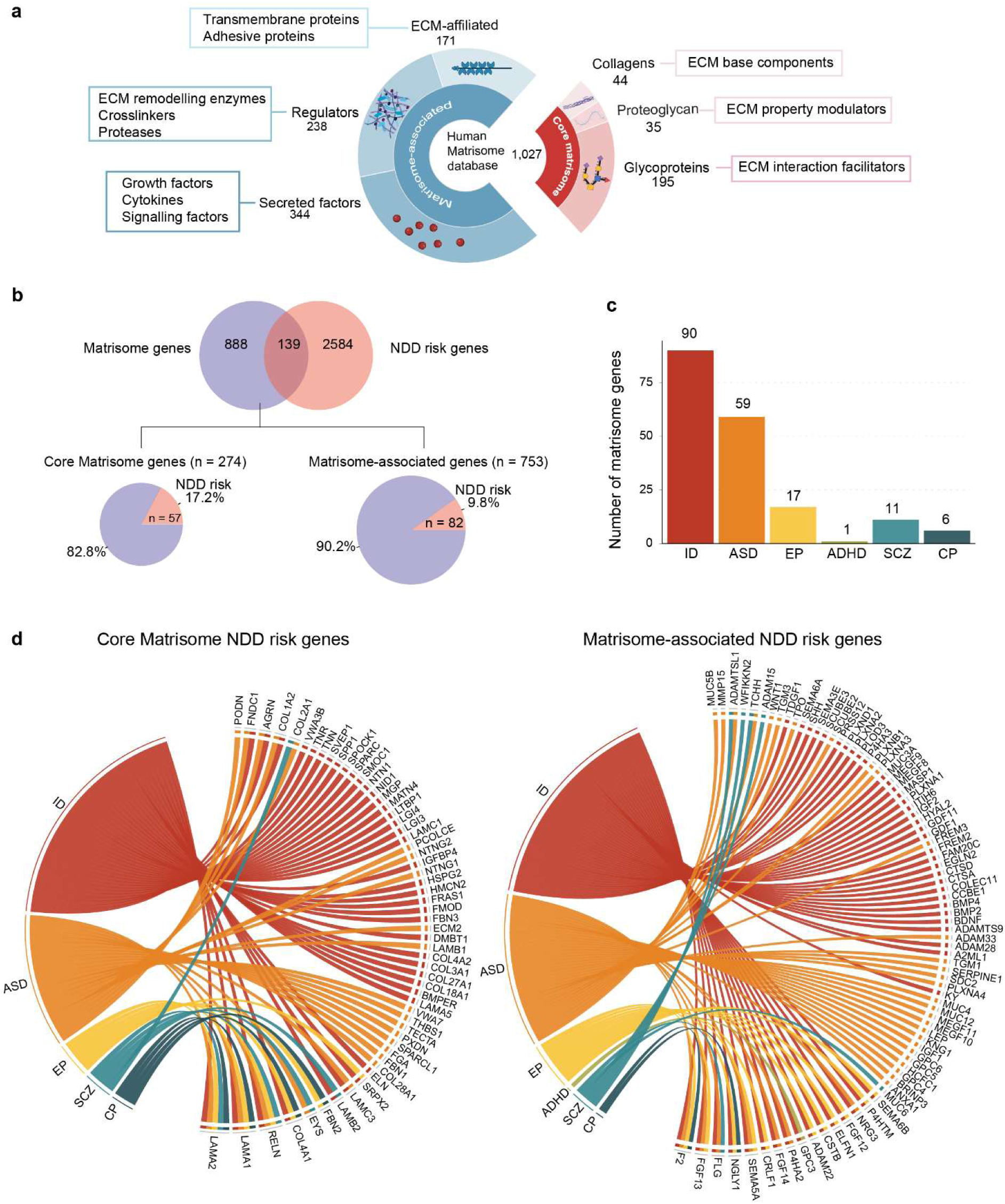
Identification of NDD-Associated Matrisome Genes. **a** Schematic overview of the different components of the core matrisome and matrisome-associated proteins. Illustration created using BioRender (BioRender.com). **b** Venn diagram illustrating the overlap between matrisome genes and NDD risk genes. Pie charts represent the proportion of core matrisome and matrisome-associated genes that are classified as NDD risk genes. **c** Number of matrisome genes identified as risk factors for each NDD. Intellectual disability (ID), autism spectrum disorder (ASD), epilepsy (EP), attention deficit hyperactivity disorder (ADHD), schizophrenia (SCZ), and cerebral palsy (CP). **d** Chord plotS showing association between NDD types and their corresponding core matrisome (left) and matrisome-associated NDD risk genes (right).

### A framework for scRNA-seq meta-analysis

To investigate cell-type-specific matrisome signatures during cortical development, we performed a comprehensive meta-analysis of scRNA-seq data from six independent studies encompassing 37 fetal cortex samples. The analysis pipeline consisted of three key steps: (1), raw count matrices were retrieved from all studies. (2), rigorous quality control of each dataset was performed by removing low-quality cells and doublets, followed by normalization and log-transformation of gene counts. (3), the datasets were integrated using 2,000 anchor genes to create a unified meta-dataset **(Fig. 2a)**. Subsequently, k-shared nearest neighbor (k-SNN) clustering on the integrated data revealed 40 distinct clusters. Each cluster was annotated with a cell type label using a semi-supervised approach that combined the scType algorithm^38,39^ with known cell type markers **(Supplementary Fig. 1a and Supplementary Table 1)**. The final integrated dataset comprised 213,659 cells spanning GW 8 to 26, with varying contributions from each study **(Fig. 2b)**. The successful integration of multiple datasets was visualized by the Uniform Manifold Approximation and Projection (UMAP), indicating minimal batch effects **(Fig. 2c)**. We checked the data set integration for robustness using two measures: (1) the integration Local Inverse Simpson’s Index (iLISI) and (2) cell-type LISI (cLISI) scores, which measure the batch-mixing and cell type grouping respectively^40^. In an ideal setting, iLISI should be close to 2, while cLISI should be close to 1. Indeed, in our integrated data iLISI was 2.004 and cLISI was 1.277 **(Fig. 2c, d)**. For comparison, we performed the same integration on a control dataset of human peripheral immune cells, achieving a similar integration with an iLISI of 1.508 and a cLISI of 1.146 **(Supplementary Fig. 1b)**. Additionally, we calculated the LISI score from the complete batch mixing of our meta-dataset, which was 2.714 **(Supplementary Fig.1c)**. These analyses further validated the robustness of our integration approach.

**Fig. 2:**
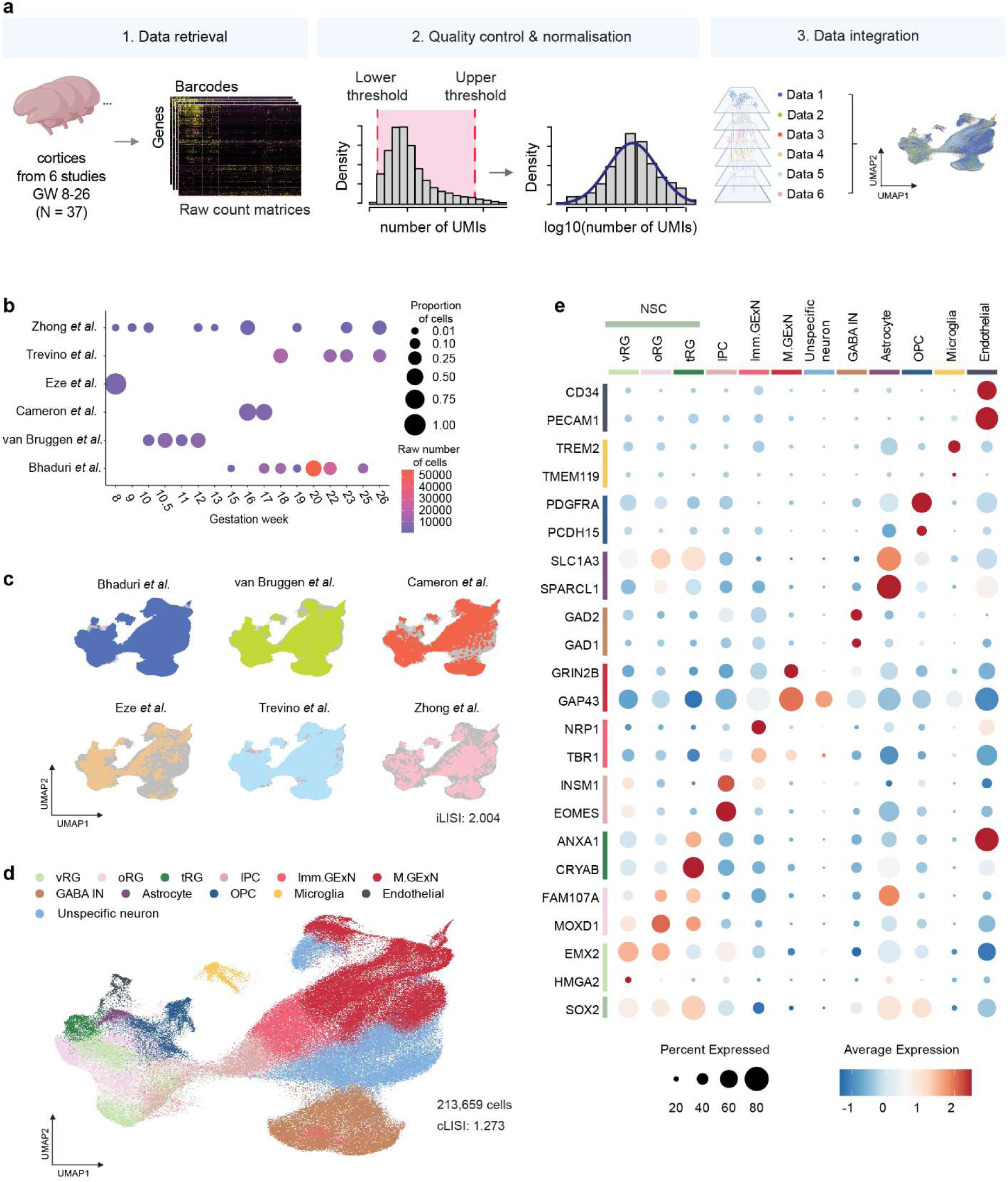
Framework of scRNA-seq meta-analysis integrating multiple datasets. **a** Overview of the processing pipeline for scRNA-seq meta-data. Illustration created using BioRender (BioRender.com). **b** Dot blot representation of each study’s contribution to the meta-dataset across different gestational weeks. The proportion of cells is indicated by dot size, while the raw number of cells is represented by a color scale. **c** Distribution of scRNA-seq data from each study on the meta-data UMAP. The integration Local Inverse Simpson’s Index (iLISI) for the meta-dataset is 2.004. **d** UMAP visualization of the integrated and clustered dataset, annotated and color-coded by 12 cell types. The cell type Local Inverse Simpson’s Index (cLISI) for the meta-dataset is 1.273. **e** Dot plot representation of the expression of canonical markers for each cell type. Dot size indicates the percentage of cells expressing the gene, while dot color represents the average expression level.

We annotated clusters into major cell types, including neural stem cells (NSC), intermediate progenitor cells (IPC), immature glutamatergic excitatory neurons (Imm.GExN), maturing glutamatergic neurons (M.GExN), unspecific neurons, GABAergic inhibitory neurons (GABA IN), astrocytes, oligodendrocyte precursor cells (OPC), as well as non-neural cell types such as microglia and endothelial cells . This analysis also revealed distinct clusters of NSC subtypes including ventricular radial glia (vRG), outer radial glia (oRG) and truncated radial glia (tRG). **(Fig. 2d, Supplementary Fig. 1a and Supplementary Table 2)**. The final annotated meta-dataset showed a balanced representation of major cell types, with each type being represented by multiple donors **(Supplementary Fig. 1d)**.

We validated cell type annotations by examining the expression levels of cell-type markers across annotated clusters **(Fig. 2e)** and inspecting the overlay of individual cell types along with the expression levels of their corresponding cell type markers on the integrated UMAP **(Supplementary Fig. 2a)**. Notably, we identified multiple clusters concordant with vRG and oRG, indicating gene expression heterogeneity within these NSC populations **(Supplementary Fig. 2b)**. We also identified a cluster of neurons that did not fit into known neuronal cell types (labeled as unspecific neuron). These neurons exhibited upregulated neuronal hemoglobin genes and increased metabolic activity, indicative of a less differentiated state, based on differential gene expression (DGE) analysis and subsequent gene ontology (GO) analyses **(Supplementary Fig. 2c-e)**.

### Cell type specific signatures of matrisome genes in the developing human cortex

We next sought to understand the cell-type-specific and shared patterns of matrisome gene expression in our integrated meta-dataset. To identify potential patterns in matrisome gene expression across cell types, hierarchical clustering was applied to a matrix of average gene expression levels per cell type. This analysis revealed the presence of distinct matrisome gene expression signatures for each cell type among 953 detected matrisome genes **(Supplementary Fig. 3)**.

To systematically identify matrisome marker genes across cell types, we performed DGE analysis using pseudobulk gene expression as input, comparing each cell type to all others. Pseudobulk aggregates gene expression data by cell type within each donor, reducing noise in the integrated meta-dataset and enhancing the biological interpretability of the analysis **(Fig. 3a)**. Most cell types exhibited distinct matrisome marker genes, with endothelial cells and astrocytes having the highest numbers (102 and 100, respectively), highlighting their prominent roles in ECM composition and remodeling as well as the propagation of signaling molecules **(Fig. 3b)**. Microglia are known to interact with various cell types and are finely tuned to adapt to environmental cues, enabling them to adjust their roles in neuroprotection, immune defense, and tissue remodeling^41^. In agreement, the unique enrichment of matrisome-associated marker genes in microglia suggests their essential contribution to these functions.

**Fig.3:**
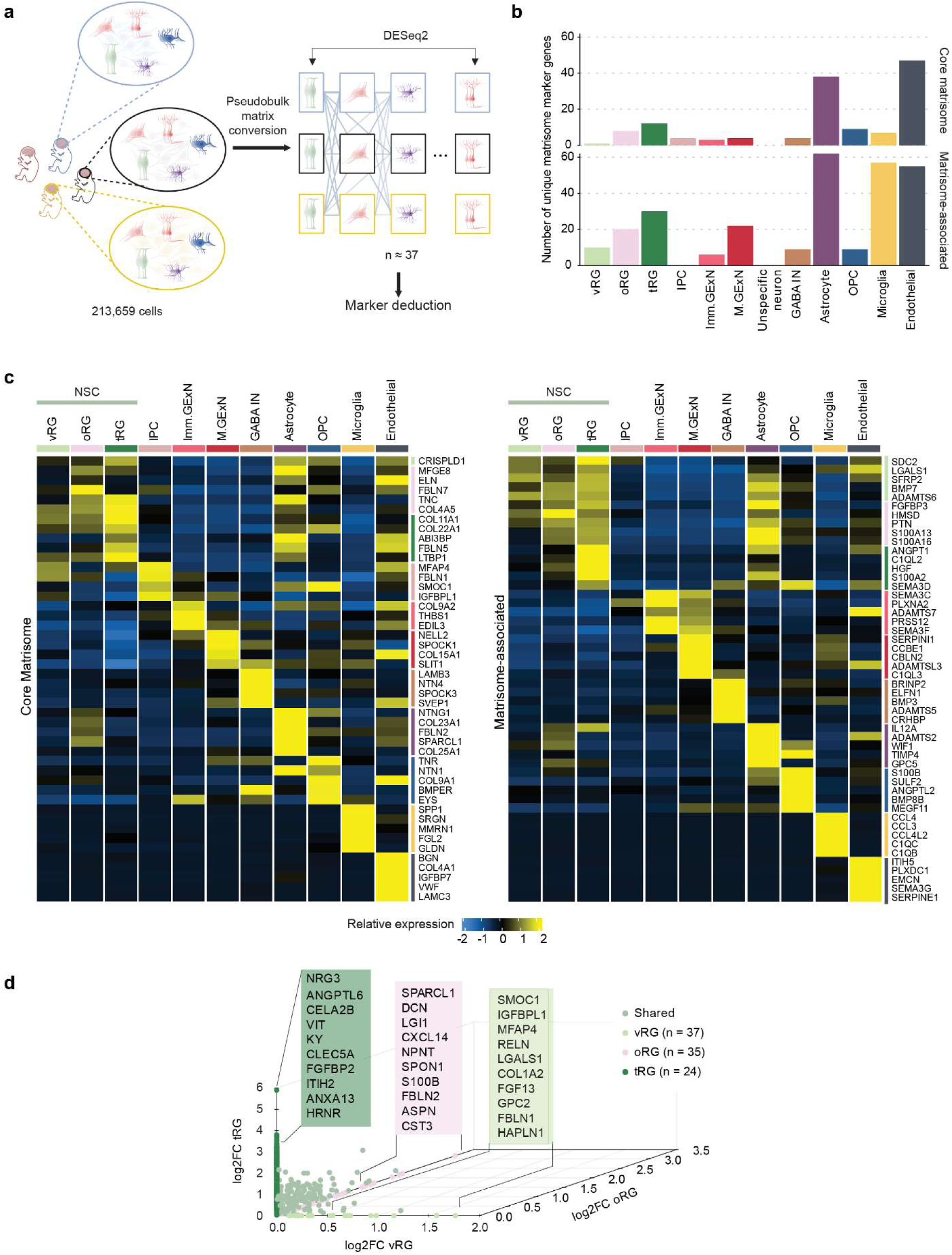
Cell type-specific matrisome signature during cortical development. **a** Schematic representation of the pseudobulk approach, illustrating the aggregation of gene expression data by cell type for each donor. Illustration created using BioRender (BioRender.com). **b** Bar graph displaying the number of unique core matrisome and matrisome-associated marker genes (log₂ fold-change > 1, adjusted p-value < 0.05) for each cell type. **c** Heatmap depicting the average expression levels of the top unique core matrisome and matrisome-associated marker genes for each cell type (ranked by log₂ fold-change, adjusted p-value < 0.05). **d** 3D scatterplot of differentially expressed matrisome genes based on log₂ fold-change among NSC subtypes: ventricular radial glia (vRG, x-axis, pale green), outer radial glia (oRG, z-axis, pink), and truncated radial glia (tRG, y-axis, dark green). The number of donors per cell type is indicated. Matrisome genes that are highly expressed in at least two NSC subtypes are highlighted in pastel green.

To further assess the specificity of cell-type matrisome marker genes, the expression levels of up to the top five core matrisome and matrisome-associated marker genes for each cell type were visualized **(Fig. 3c and Supplementary Table 3)**. The heatmaps revealed distinct matrisome gene signatures for each cell types. While some genes, such as *SPP1* in microglia and *SEMA* family genes in neuronal cells, are known to be enriched in these cell types, many remain unexplored in the context of cortical development, suggesting novel targets for future functional studies. Collagen, a key ECM component in the developing cortex, plays a crucial role in structural organization, neural stem cell behavior, neuronal migration, vascular development, and intercellular signaling^42^. Notably, our findings highlight that each cell type exhibits preferential expression of specific collagen subtypes **(Fig. 3c)**. We observed that NSC subtypes share a common expression pattern of matrisome marker genes **(Fig. 3c)**. To identify the distinct features of matrisome gene expression in each NSC subtype, we conducted additional DGE analysis comparing matrisome gene expression among vRG, oRG, and tRG **(Fig. 3d, Supplementary Tabel 4)**. The differentially expressed matrisome genes in each NSC subtype reflect their unique functional roles in cortical development. In vRG, matrisome genes such as *RELN*, *COL1A2*, and *SMOC1,* which support ECM remodeling, neuronal migration, and structural organization^43–45^, were upregulated. In oRG, *SPARCL1*, *LGI1*, *FBLN2* and *S100B*^46–48^ showed relatively higher expression, contributing synaptic connectivity and organization, neurogenesis, and gliogenesis. In tRG, genes such as *NRG3*, *FGFBP2*, and *ITIH2* were enriched, facilitating ECM remodeling, signaling, and adaptive progenitor activity^49–51^. These distinct matrisome expression profiles suggest specialized functions of each NSC subtype during cortical development **(Fig. 3d)**.

Collectively, our analyses identified unique, cell-type-specific matrisome signatures, suggesting the potential distinct roles of matrisome components in supporting each cell type’s specialized functions during cortical development.

### Temporal dynamics of matrisome gene regulation during cortical development

Cortical development follows distinct temporal dynamics, characterized by shifts in cellular behavior, composition, and cytoarchitecture. NSCs transition from a proliferative to a neurogenic state, followed by gliogenesis, while the cortex undergoes layer formation, neuronal migration, and structural maturation, ultimately establishing its complex architecture^52^.

To examine temporal changes in matrisome gene expression across all cell types during cortical development, particularly in NSCs, given their dynamic behavior throughout the developmental trajectory, we analyzed matrisome expression across three periods: late first trimester (GW 8– 12), early-second trimester (GW 13–19), and late-second trimester (GW 20–26). We used a linear model to evaluate the relationship between gene expression in each cell type across all three developmental periods. The analysis revealed distinct temporal dynamics of matrisome gene expression across cell types in the developing human cortex (**Fig. 4a and Supplementary Table 5**). NSCs show notable increases in *TNC*, *LGALS3*, and *TIMP3*, while *FBLN1* exhibited a decreasing trend. Neuronal cell types demonstrate unique matrisome signatures, with *MDK* showing a pronounced increase in IPC and M.GExN, and *PXDN* showing a temporal decrease in Imm.GExN and GABA IN. This signature is consistent with the previous understanding of *MDK*’s role in neural plasticity^53^ and *PXDN*’s function in proliferation^54^. The analysis also revealed that all cell types display a higher number of temporally increasing matrisome genes than temporally decreasing genes **(Supplementary Table 5)**. Non-neuronal lineage cell types exhibit the strongest tendency toward positive temporal regulation, with endothelial cells showing an 85.0% (17 out of 20 genes) and OPCs displaying 80.6% (25 out of 31 genes) of temporally increasing matrisome genes, suggesting their growing contribution to matrisome-mediated functions during cortical development (**Supplementary Table 5**).

**Fig. 4:**
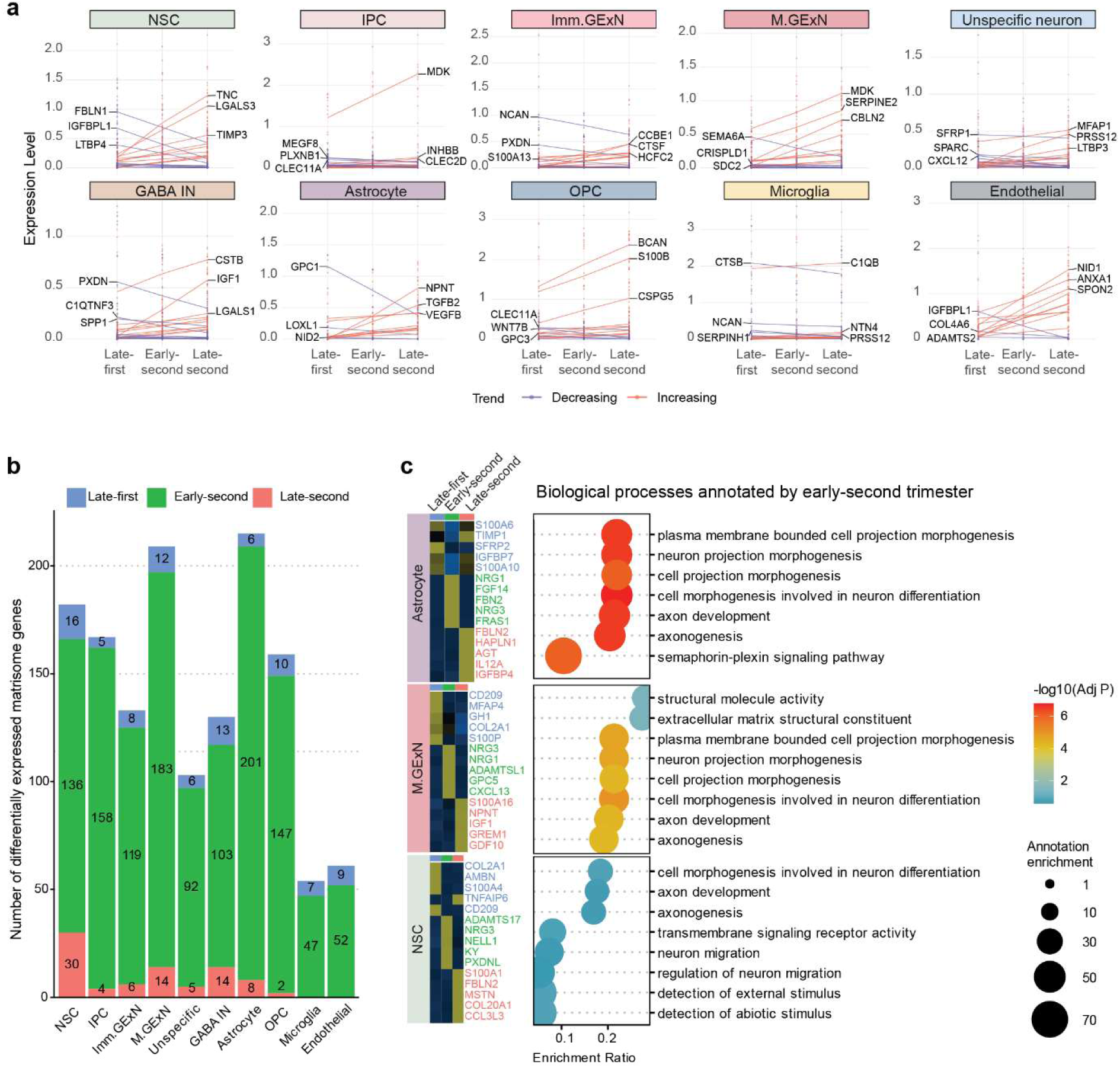
Temporal dynamic signatures of matrisome genes in each cell type during cortical development. **a** Temporal expression patterns of matrisome genes across distinct cell types during human cortical development. Each data point represents the expression level of matrisome genes in an individual donor, with a best-fitted trend line illustrating the expression pattern. Expression levels of the top 10 matrisome genes showing temporal increases (red) and decreases (blue) are displayed, and the three most significant genes are labeled for each cell type across three developmental periods. **b** Bar graph displaying the number of differentially expressed matrisome genes in each cell type across three developmental periods (log₂ fold-change > 1, adjusted p-value < 0.05). **c** On the left, a heatmap displaying the expression levels of up to the top five matrisome genes across three developmental periods in astrocyte, GExN, and NSC (ranked by log₂ fold-change, adjusted p-value < 0.05). On the right, GO enrichment analysis of early-second trimester marker genes (ranked by log₂ fold-change, adjusted p-value < 0.05) against a matrisome gene background in astrocyte, GExN, and NSC. The x-axis represents the enrichment score, dot size indicates the number of genes in the annotation, and the -log₁₀ adjusted p-value is represented by the color scale

Next, we sought to identify matrisome gene expression patterns specific to a particular development period in each cell type. We analyzed cell-type-specific temporal matrisome expression signatures using DGE analysis across three developmental periods. The early-second trimester exhibited a significantly higher number of differentially expressed matrisome marker genes **(Fig. 4b and Supplementary Table 6)**. This reflects robust matrisome activity during this stage, aligning with the diverse cellular processes occurring at this period. We visualized up to the top five matrisome marker genes for each developmental stage in each cell type, revealing distinct and stage-specific temporal matrisome expression signatures (**Supplementary Fig. 4 and Supplementary Table 6**). Astrocytes, M.GExN, and NSCs had the highest number of temporal marker genes, suggesting significant contribution of matrisome gene expression during development. To explore the functions of their matrisome genes during the early-second trimester, GO enrichment analysis was performed on the early-second trimester matrisome marker genes for those cell types. In astrocytes and M.GExN, genes associated with cell morphogenesis, including *FBN2*, *FRAS1*, *ADAMTSL1*, and *GPC5*, as well as genes involved in axon development, such as *NRG1*, *NRG3*, and *FGF14*, were enriched. In NSCs, genes involved in axon development, neuronal migration and maturation, including *NRG3*, *ADAMTS17*, *NELL1*, *KY*, and *PXDNL* were enriched **(Fig. 4c)**. These represent important functions of temporally fine-tuned expression of matrisome genes in each cell type during development. In particular, the enriched expression of matrisome genes associated with axon development, cellular morphogenesis, and neuronal maturation and migration in astrocytes and NSCs highlights the importance of cell-to-cell interactions (CCI) mediated by the matrisome.

### Matrisome-mediated cell-cell communications during cortical development

To examine the role of matrisome genes in cell communications, we analyzed CCI and signaling pathways mediated by matrisome genes using CellChat^55^. Analysis of cell-type-specific interaction networks revealed distinct differences between whole-transcriptome-wide and matrisome-specific communication in the developing cortex. The whole transcriptome-based network showed dense interconnectivity across all cell types, with strong interactions primarily involving glial cells (vRG, oRG, tRG, OPC, and astrocytes) and endothelial cells. In contrast, the matrisome-specific network exhibited selective connectivity, with tRG, astrocytes, and endothelial cells emerging as major contributors to CCI. Notably, interactions among tRG, astrocytes, and endothelial cells in the whole transcriptome heavily rely on matrisome genes. Additionally, autocrine interactions mediated by matrisome genes were exclusively observed in tRG and astrocytes **(Fig. 5a)**. These analyses indicate that the matrisome may play a role in communication between specific cell types during cortical development.

**Fig. 5:**
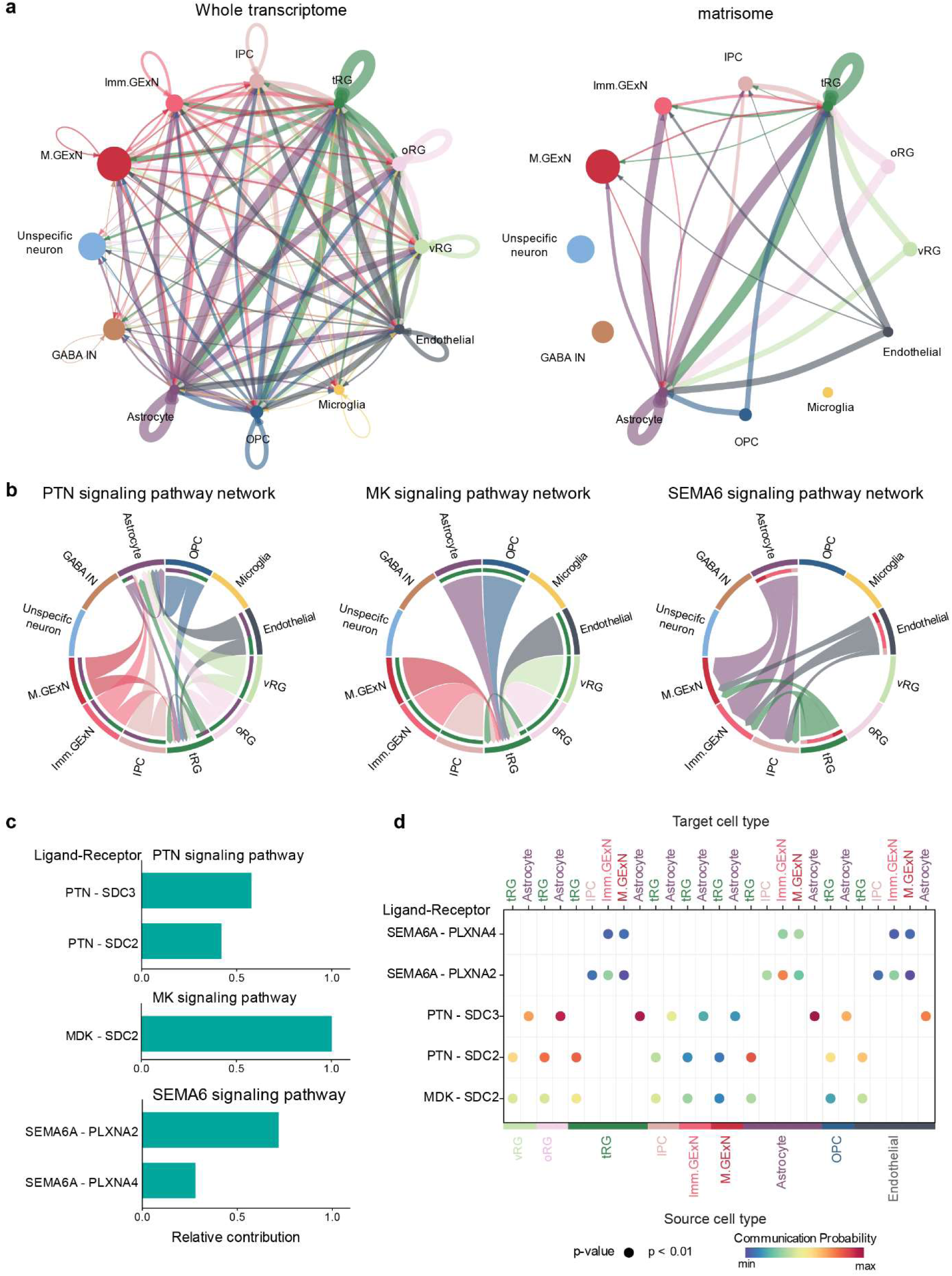
Cell-cell communication mediated by matrisome. **a** Network visualization of interaction weights/strength between cell types based on whole transcriptome (left) and matrisome (right) gene expression. Node size represents the relative abundance of cell types in the meta-data, while edge thickness indicates interaction strength. Arrow direction denotes the flow of signaling from sender to receiver. **b** Chord plots illustrating cell-type-specific signaling networks for PTN, MK, and SEMA6 pathways. Arc width represents the interaction strength between cell types. **c** Bar graph showing the relative contribution of individual ligand-receptor pairs to overall pathway activity. Bar length represents the normalized contribution strength **d** Dot plot illustrating source-target cell type relationships for each ligand-receptor pair. Dot size represents statistical significance (p < 0.01), while color intensity indicates communication probability.

Signaling pathway-specific analysis identified three matrisome-mediated signaling networks during cortical development: the pleiotrophin (PTN), midkine (MK), and semaphorin 6A (SEMA6) pathways **(Fig. 5b)**. The PTN pathway, known for its role in modulating proliferation, differentiation, and cell survival^56,57^, exhibited robust bidirectional communication between neural and glial cells, with signaling notably converging on tRG and astrocytes. The MK signaling network, which supports cell growth and neural plasticity^58^, showed a more focused interaction pattern, primarily targeting tRG. In contrast, SEMA6 signaling, associated with neuronal morphogenesis and migration^59,60^, revealed distinct interactions targeting neuronal populations. Notably, tRG, astrocytes, and endothelial cells were identified as source cell types for all three pathways, highlighting their active roles in coordinating cell communication during cortical development **(Fig. 5b)**.

Quantitative analysis of ligand-receptor pairs identified PTN-SDC3, MDK-SDC2, and SEMA6A-PLXNA4 as the key contributors within their respective signaling pathways **(Fig. 5c)**. This analysis also uncovered cell-type-specific communication patterns, with significant interactions observed between distinct cellular populations. Notable examples include strong PTN-SDC3 signaling from vRG/oRG to tRG and astrocytes, as well as SEMA6A-PLXNA4 signaling from astrocytes to Imm.GExN and M.GExN **(Fig. 5d)**.

Together, these findings highlight the matrisome’s specialized role in mediating cell-cell communication during cortical development, with tRG, astrocytes, and endothelial cells emerging as key potential regulators of these interactions.

### Characterization of neuronal lineage specific changes in matrisome gene expression

NSCs exhibit substantial heterogeneity during cortical development, shaped by their diverse fate choices along distinct developmental trajectories^33^. To explore matrisome signatures during neurogenesis, we performed DGE analysis to compare NSCs, IPCs, and excitatory neurons (GExN, including Imm. GExN and M.GExN). This analysis revealed distinct changes in matrisome signatures along the trajectory of the neurogenic lineage (**Fig. 6a and Supplementary Table 7)**. Among the differentially expressed matrisome genes, *LGALS3* stood out due to its pronounced temporal dynamics in NSCs **(Fig. 4a)** and a significant expression difference between NSCs and GExN (log2 fold-change > 4). *LGALS3* encodes β-galactoside-binding Galectin-3 (GAL3) and has been suggested as a transcriptional marker for oRG through scRNA-seq^61^, but its validation has not been previously reported. Consistent with these findings, weighted gene co-expression network analysis (WGCNA) and Pearson’s correlation analysis revealed that *LGALS3* is co-expressed with *HOPX*, a well-established oRG marker **(Fig. 6b, c)**. To validate the expression of *LGALS3* and *HOPX*, we performed immunofluorescent co-staining of GAL3 and HOPX in human fetal prefrontal cortices at GW16 and GW17. 71.1 ± 4.2% of HOPX^+^ cells expressed GAL3 **(Fig. 6d-e)**. The proportion of cells with high *LGALS3* expression increased during the early-second trimester (GW13–19), which was confirmed by immunofluorescence staining **(Fig. 6f, g)**.

**Fig. 6:**
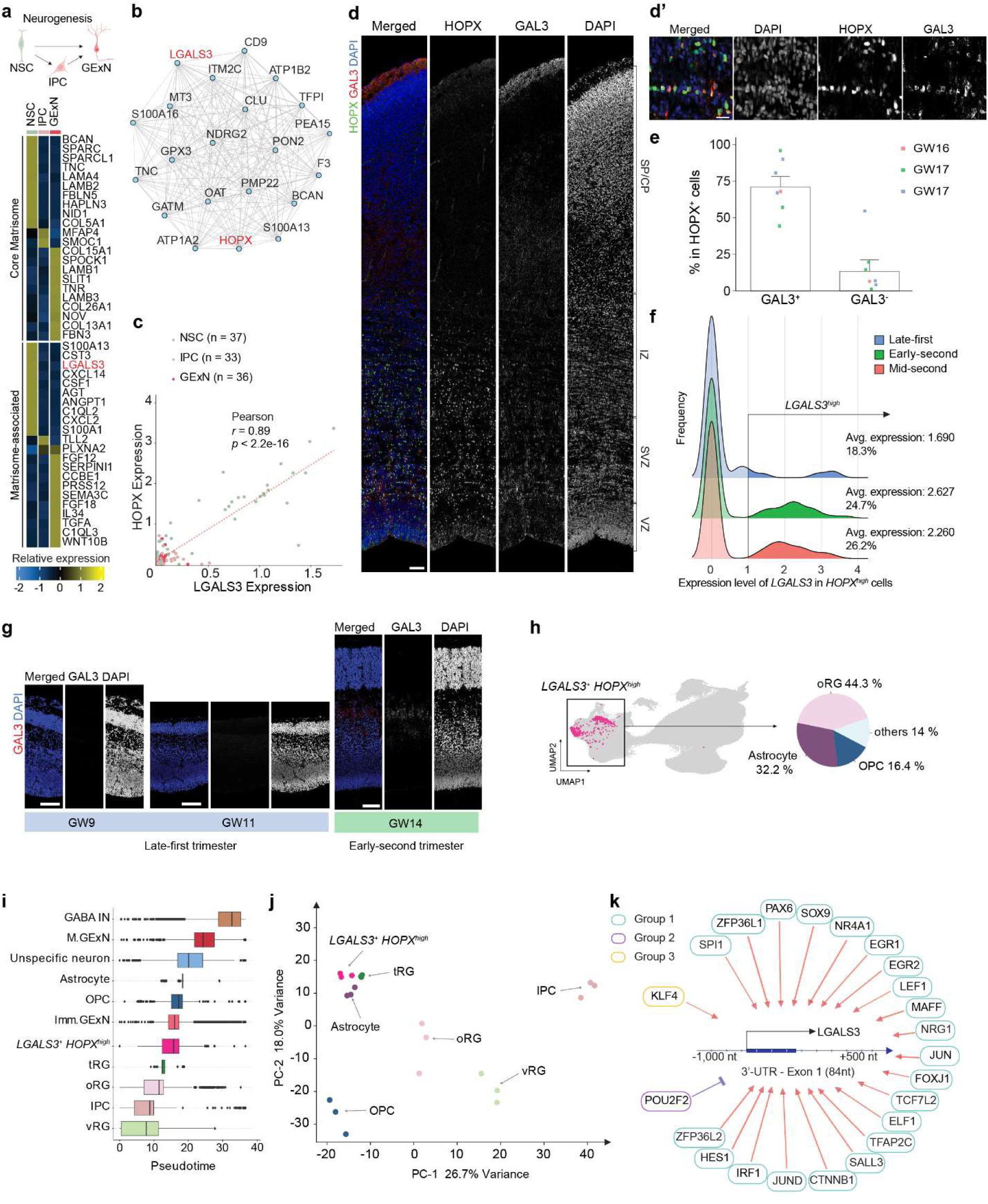
Characterization of neuronal lineage specific changes in matrisome gene expression. **a** Illustration depicting the neurogenic lineage, created using BioRender (BioRender.com). Heatmap displaying the expression levels of up to the top 10 cell-type-specific core matrisome (top) and matrisome-associated (bottom) marker genes across cells in the neurogenic lineage (ranked by log₂ fold-change, adjusted p-value < 0.05). **b** Co-expression network of LGALS3 across NSC, IPC, and GExN, highlighting the top significantly co-expressed genes identified through weighted gene co-expression network analysis. **c** Scatter plot showing the correlation between LGALS3 and HOPX in scRNA-seq meta-data. The Pearson correlation coefficient and p-value for the best-fitted line are displayed. Each colored outline represents a donor per cell type. **d-d’** Immunofluorescence staining of HOPX, Galectin-3 (GAL3), and DAPI in the GW16 fetal prefrontal cortex. The inset provides a magnified view. Scale bars: 100 μm and 10 μm (inset) **e** Quantification of GAL3⁺HOPX⁺ and GAL3⁻HOPX⁺ cells as a percentage of the total HOPX⁺ cell population from immunofluorescence staining in human fetal prefrontal cortices at GW16–17. Data are from three donors and presented as mean ± SEM. **f** Density plots illustrating the frequency distribution of HOPX^high^ NSCs with LGALS3 expression across human fetal developmental stages. LGALS3^high^ is defined as an expression level >1, as indicated on the horizontal axis **g** Immunofluorescence staining GAL3 and DAPI in the fetal prefrontal cortex at GW 9, 11, and 14. Scale bars: 100μm. **h** UMAP of scRNA-seq meta-data highlighting *LGALS3^+^HOPX^high^*cells (magenta) with a pie chart showing the cellular composition of *LGALS3^+^HOPX^high^*population **i** Box plots illustrating the developmental trajectory across different cell types in the scRNA-seq meta-data. The x-axis represents pseudotime values, with boxes indicating the median and quartiles of each population’s distribution. Cell populations are arranged in descending order from the top based on their mean pseudotime. **j** Principal component analysis plot illustrating the transcriptional relationships between different cell populations. **k** Schematic representation of the LGALS3 regulatory region and its associated transcription factors (TFs). The diagram illustrates the genomic region spanning -1,000 to +500 nucleotides relative to the LGALS3 transcription start site. Red arrows indicate transcriptional activation, while blue blunt arrow represents transcriptional inhibition. Each group represents a module of TFs with a shared expression pattern over pseudotime.

While oRG shows strong *HOPX* expression, vRG exhibits minimal expression^61^. To examine the potential role of *LGALS3* in oRG, we further identified *LGALS3^+^* cells with high *HOPX* expression (*HOPX^high^*). More than 44% of *LGALS3^+^HOPX^high^* were annotated as oRG. Additionally, a significant proportion of *LGALS3*^+^*HOPX^high^* were annotated as macroglia, including astrocytes and OPCs, potentially indicating their differentiation into macroglia **(Fig. 6h)**. To explore the developmental positioning of *LGALS3^+^HOPX^high^* cells along a differentiation trajectory, we performed pseudotime analysis with vRG as the root cell type **(Supplementary Fig. 5a)**. The analysis showed that *LGALS3^+^HOPX^high^*cells were assigned a higher pseudotime value, consistent with a more mature differentiation state compared to vRG, oRG, tRG, and IPC **(Fig. 6i).** To understand the transcriptional similarities among *LGALS3^+^HOPX^high^*cells and other cell types, we performed principal component analysis (PCA) on the pseudobulk gene expression profile. Prior to the analysis, we randomly assigned cells into three arbitrary groups and aggregated gene expression data to mitigate potential noise from gestational variability among donors and the large cell numbers in each cell type in our integrated dataset. PCA showed *LGALS3^+^HOPX^high^*cells clustering near astrocyte and tRG, distinct from IPC and OPC **(Fig. 6j)**. Our analyses suggest that *LGALS3^+^HOPX^high^* cells may represent a subpopulation of NSCs with a preferential lineage trajectory towards astrocytes.

To further examine the transcriptional regulators of *LGALS3+* cells, we conducted an integrative meta-analysis utilizing ATAC-seq and ChIP-seq datasets from the ChIP-Atlas database^62–64^. Our analysis revealed that *LGALS3* promoter region and transcription start site (TSS) are predominantly in an open chromatin state in the human frontal cortex **(Supplementary Fig. 5b)**. Notably, H3K27me3 marks, enriched in neural progenitor cells suggest a potential primed transcriptional state, while astrocytes exhibit H3K27ac marks, indicating active transcription **(Supplementary Fig. 5b)**^65,66^. We next performed integrated regulatory network analysis (IReNA) to identify transcription factors (TFs) that bind to the promoter regions of *LGALS3* potentially regulating its expression^67^. Twenty-three TFs were identified and grouped by shared expression patterns along the developmental trajectory, exhibiting either positive or negative co-expression with *LGALS3*. Importantly, the genomic region spanning 1,000 nucleotides upstream to 500 downstream of *LGALS3*’s TSS contains binding motifs for these TFs, suggesting potential direct regulatory roles **(Fig. 6k and Supplementary Table 8)**. GO analysis revealed that transcriptional activators of *LGALS3* are associated with gliogenesis and glial cell differentiation (**Supplementary Fig. 5c**). Collectively, these findings suggest an astrocytic developmental trajectory for *LGALS3^+^*cells.

### Characterization of macroglial lineage specific changes in matrisome gene expression

Gliogenesis, alongside neurogenesis, is a key lineage specification process in the developing cortex. To investigate matrisome gene expression dynamics during gliogenic lineage progression, we analyzed the matrisome signatures of NSCs, astrocytes, and OPCs using DGE analysis. The heatmap revealed distinct expression patterns of core and matrisome-associated marker genes across NSC, astrocyte, and OPC populations **(Fig. 7a and Supplementary Table 9)**. Astrocytes and OPCs exhibited a broader range of matrisome markers than NSCs. Notably, *S100B*, a well-established astrocyte marker, was enriched in OPCs. While *S100B* is known to be expressed in OPCs and immature oligodendrocytes in the developing mouse brain^68,69^, its expression dynamics in OPCs during human cortical development remain poorly understood.

**Fig. 7:**
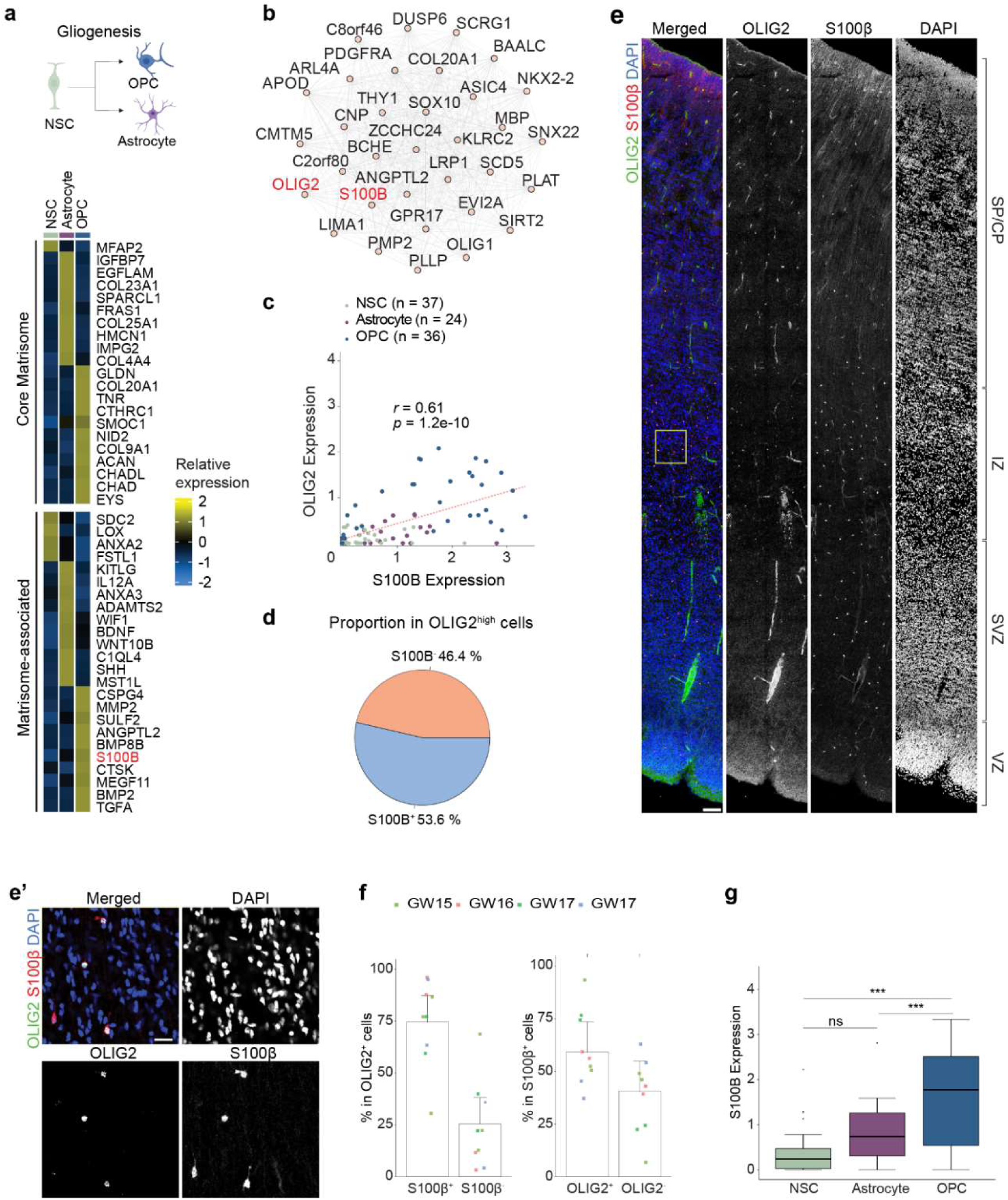
Characterization of macroglial lineage specific changes in matrisome gene expression. **a** Illustration depicting the macroglial lineage, created using BioRender (BioRender.com). Heatmap displaying the expression levels of up to the top 10 cell-type-specific core matrisome (top) and matrisome-associated (bottom) marker genes across cells in the macroglial lineage (ranked by log₂ fold-change, adjusted p-value < 0.05). **b** Co-expression network of OLIG2 across NSC, astrocyte, and OPC, highlighting the top significantly co-expressed genes identified through weighted gene co-expression network analysis (WGCNA). **c** Scatter plot showing the correlation between S100B and OLIG2 in scRNA-seq meta-data. The Pearson correlation coefficient and p-value for the best-fitted line are displayed. Each colored outline represents a donor per cell type. **d** Pie chart showing the proportion of S100B^+^ cells among OLIG2^high^ cells in the scRNA-seq meta-data **e-e’** Immunofluorescence staining of OLIG2, S100B, and DAPI in the GW16 fetal prefrontal cortex. The inset provides a magnified view. Scale bars: 100 μm and 10 μm (inset) **f** Quantification of S100β^+^ OLIG2^+^ and S100β^-^ OLIG2⁺ cells as a percentage of the total OLIG2⁺ cell population (left) and S100β^+^ OLIG2^+^ and S100β^+^ OLIG2^-^ cells as a percentage of the total S100β⁺ (right) from immunofluorescence staining in human fetal prefrontal cortices at GW15–17. All data are from four donors and presented as mean ± SEM. **g** Box plots showing the mean expression levels of *S100B* in NSCs (n = 37), astrocytes (n = 4), and OPCs (n = 36) in the scRNA-seq meta-data. The y-axis represents the expression level of S100B, with boxes indicating the median and quartiles of each population’s distribution. One-way AONVA, p = 7.1e^-6^, ***: p<0.001, ns: not significantly different.

WGCNA revealed significant co-expression of *S100B* with *PDGFRA*, *OLIG1*, and *OLIG2* **(Fig. 7b)**. Moreover, a strong correlation between *OLIG2* and *S100B* expression in OPCs was confirmed by Pearson’s correlation analysis (coefficient: 0.61, **Fig. 7c**), with 53.6% of *OLIG2^high^* cells expressing *S100B* **(Fig. 7d)**. To validate this, immunofluorescence staining of S100β and OLIG2 was performed in human fetal prefrontal cortices at GW15 and GW17, revealing that 74.8 ± 6.4% of OLIG2^+^ cells are S100β^+^ **(Fig. 7e-f)**. To assess S100B expression patterns in the context of fate specification, we performed a comparative analysis of mean expression levels of each donor for NSCs, astrocytes and OPCs, revealing that OPCs exhibit significantly higher *S100B* expression levels than NSCs or astrocytes **(Fig. 7g)**. Collectively, our analyses identify the strong expression of S100B in OPCs during human cortical development.

### Cell type and temporal-specific expression of matrisome genes associated with NDDs

A fundamental approach to understanding the pathogenesis of NDDs is to study the function of NDD risk genes. A major challenge lies in examining gene function within the appropriate cell-type and developmental context. Through our analyses, we identified cell-type- and temporal-specific signatures of matrisome genes in the developing cortex **(Figs.3-4)**. Accordingly, we sought to characterize cell type specific and time windows of NDD risk matrisome gene expression, providing insights for the design of functional studies of risk genes. First, we quantified the number of cell-type-specific matrisome marker genes associated with specific NDDs across different cell types. The analysis revealed that matrisome marker genes of tRG, astrocytes, OPCs, and endothelial cells show the highest number of matrisome marker genes associated with NDDs **(Fig. 8a and Supplementary Table 10)**. ID exhibited the highest number of enriched matrisome marker genes across all cell types, with a particularly strong representation in endothelial cells. In contrast, matrisome marker genes associated with ASD were predominantly enriched in non-neural cells. To assess the specificity of cell-type matrisome marker genes associated with NDDs, the expression levels of the top matrisome marker genes were analyzed and visualized in volcano plots and a heatmap **(Fig. 8b-c, Supplementary Fig. 6 and Supplementary Table 11)**. Some NDD risk matrisome genes exhibited high cell-type specificity, with well-characterized functional significance. For example, *SPP1*, an ID risk gene, is highly expressed in microglia and was recently identified as a key regulator of structural integrity during brain development in mice^70^. *COL4A1*, predominantly expressed in endothelial cells, is linked to multiple NDDs, including ID, EP, CP, and neonatal hemorrhages, and plays a crucial role in human fetal vascular development^71^ **(Fig. 8c)**.

**Fig. 8:**
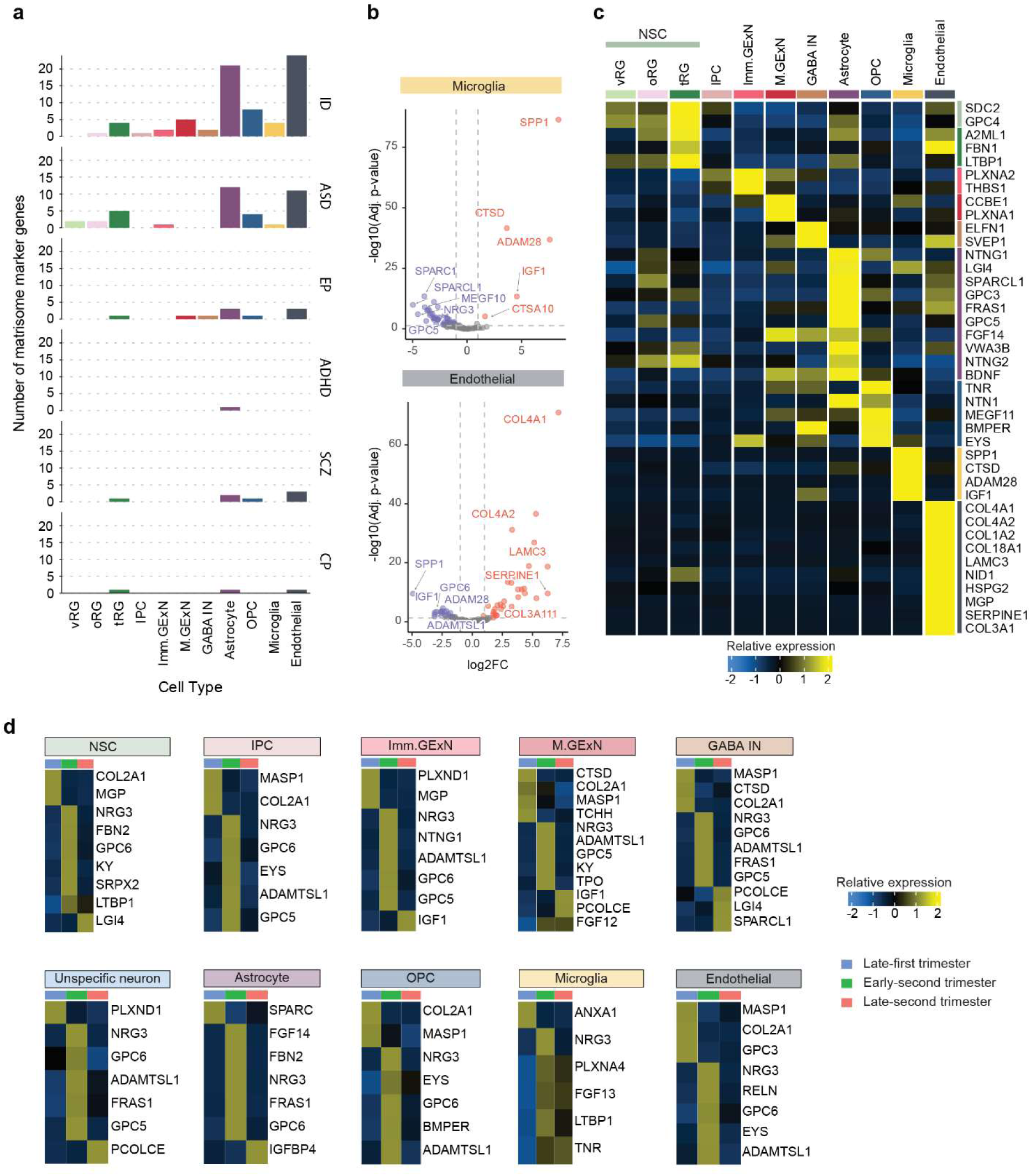
Cell type and temporal-specific expression of matrisome genes associated with NDDs. **a** Bar graphs displaying number of matrisome marker genes (y-axis) in each cell type (x-axis) associated with NDD. **b** Volcano plot illustrating differentially expressed matrisome genes in microglia and endothelial cells. The top five significantly upregulated and downregulated NDD associated matrisome genes are labeled (ranked by log₂ fold-change, adjusted p-value < 0.05). **c** Heatmap displaying the expression levels of up to the top five NDD-associated matrisome genes across different cell types (ranked by log₂ fold-change, adjusted p-value < 0.05). **d** Heatmap displaying the expression levels of up to the top five NDD associated matrisome genes across three developmental periods in all cell type (ranked by log₂ fold-change, adjusted p-value < 0.05).

Next, we investigated the temporal specificity of NDD risk matrisome genes in each cell type using DGE analysis. All cortical cell types exhibited temporally distinct NDD risk matrisome genes, particularly during the late first and early-second trimester **(Fig. 8d and Supplementary Table 12)**. Consistent with our analysis of the temporal dynamics of matrisome genes (**Fig. 4)**, many NDD risk matrisome genes were predominantly enriched in the early-second trimester. Notably, *NRG3* was enriched across all cell types during this period, aligning with its expanding roles in glial cell growth, excitatory synapse development, and neural plasticity^72,73^.

Together, our analyses identified cell-type- and temporal-specific NDD-associated matrisome genes. These findings not only highlight the importance of matrisome gene regulation in cortical development and NDD pathogenesis but also provide insights for future functional studies.

## DISCUSSION

The ECM plays a pivotal role in cortical development, acting as a structural scaffold that supports cellular organization, migration, and differentiation^2,74^. Despite its fundamental importance, the ECM’s specific contributions to cortical development and its implications for NDDs remain largely unexplored. Our study identified that a substantial portion of core matrisome genes (17.2%) and matrisome-associated genes (9.8%) are NDD risk genes, emphasizing the ECM’s critical role in cortical development **(Fig. 1b)**. While most of NDD risk matrisome genes are disease-specific, some, such as *LAMA1, LAMA2, RELN, COL4A1, SEMA5,* and *FGF13* are associated with multiple NDDs **(Fig. 1d)**. Future studies on the matrisome associated with NDD risk will provide deeper insights into the core genetic drivers that converge on common ECM-related biological pathways and mechanisms, advancing our understanding of NDD pathogenesis. *COL2A1*, a risk gene for ASD and SCZ, and *NTN1*, a risk gene for ID **(Fig. 1d)**, are notably enriched in the VZ and SVZ of the developing human cortex compared to the developing mouse cortex^25,75^. This enrichment suggests that these genes play unique roles in human brain development, potentially contributing to the complexity and specialization of the human brain. The differential expression of these genes in humans compared to mice may help explain human the susceptibility to NDDs, highlighting species-specific aspects of gene regulation and function during critical stages of cortical development.

Our study provides a comprehensive understanding of matrisome gene expression across different cell types and developmental stages, offering deeper insights into the role of the ECM in brain development. We identified distinct matrisome gene expression signatures for each cell type **(Fig. 3 and Supplementary Fig. 3)**, suggesting that cell-type-specific biological processes and functions are, in part, mediated by the matrisome.

In this study, we demonstrated that matrisome gene expression signatures exhibit temporal dynamics, with the highest number of temporally specific genes observed during the early second trimester across various cell types in the developing cortex (**Fig. 4 and Supplementary Fig. 4**). Our findings align with established research showing that the human cerebral cortex undergoes significant transcriptional dynamics and extensive cell population diversification during the second trimester^32,76^. Importantly, GO analysis revealed that these signatures are associated with key developmental processes, such as morphogenesis, migration, growth, and cell communication. This highlights the distinct and temporally regulated role of ECM components in guiding critical processes, like cellular differentiation, proliferation, and structural organization during cortical expansion^4–7,12–14,77^.

Additionally, our CCI analysis highlights that matrisome genes can exert specialized roles in cell communication in a cell type-selective manner (**Fig. 5**). Notably, non-neuronal cell types, such as tRG, astrocytes, and endothelial cells exhibited strong matrisome-mediated communication, emphasizing their crucial role during cortical development.

Furthermore, we found that matrisome gene expression signatures undergo dynamic changes during lineage specification **(Fig. 6A, 7A)**. Characterizing cells expressing these genes provides insights into the molecular mechanisms driving cellular diversity in the developing cortex. Notably, *LGALS3* emerged as one of the matrisome signature genes distinctly different from neural lineage cells, marking subpopulations of NSCs. PCA and epigenomic analyses of *LGALS3* promoter regions revealed shared transcriptomic features with astrocytes, indicating a potential correlation between *LGALS3* expression and the astrocytic lineage specification of NSCs (**Fig. 6 and Supplementary Fig. 5)**. In addition, we identified S100B, a well-known astrocyte marker as a matrisome marker gene in OPC during macroglial lineage specification. Future functional studies on these matrisome signatures will offer deeper insights into the molecular mechanism underlying cell type specification during cortical development.

Finally, we identified the cell-type-specific and temporally specific expression of matrisome marker genes associated with NDDs **(Fig. 8 and Supplementary Fig. 6)**. Understanding which matrisome genes are active in specific cell types during distinct time windows enables targeted experiments and interventions, allowing researchers to manipulate gene expression or function in relevant contexts. This information is invaluable for developing therapies targeting specific pathways or cell populations, potentially leading to more precise and effective treatments for developmental brain disorders.

The role of the matrisome during human cortical development remains largely unexplored, and our systematic analyses provide valuable initial insights. However, several limitations must be acknowledged. One limitation is our reliance on transcriptomic data, which does not fully capture the complexity of matrisome protein function, especially given the extensive post-translational modifications such as glycosylation and secretion processes that many ECM components undergo. This is particularly important, as glycosylation can significantly influence the structure, stability, and interactions of ECM proteins^78^. To address these gaps, future research should incorporate proteomics and glycomics^79^ approaches to provide a more comprehensive understanding of the matrisome. Spatial proteomics would offer critical insights into the localization and functional contexts of ECM components in their natural milieu within the developing brain. Additionally, understanding the spatial distribution of these proteins will be essential for deciphering the specific roles they play in various regions and cell types during cortical development. Integrating these methodologies will advance our knowledge of how ECM modifications contribute to normal brain development and the pathogenesis of neurodevelopmental disorders, potentially paving the way for new therapeutic strategies.

## METHODS

### Matrisome and NDD risks gene data collection

The Human matrisome database (http://matrisomeproject.mit.edu/other-resources/human-matrisome/)^80^ was used to compile a list of 1,027 matrisome genes. Neurodevelopmental risk gene lists were generated by combining data from SFARI Gene (1,162 genes, https://gene.sfari.org/), Geisinger (1183 genes, https://dbd.geisingeradmi.org/) and SysNDD (1,372 gene, https://sysndd.dbmr.unibe.ch/)^35–37^.

### scRNA-seq and snRNA-seq data collection

Count matrices from scRNA-seq and snRNA-seq datasets were retrieved from six independent studies^28–33^; Bhaduri *et al*. and Eze *et al.* accessed from The Neuroscience Multi-omic (NeMO: https://nemoarchive.org/) data Archive (RRID:SCR_002001 and RRID:SCR_002001), van Bruggen *et al*. and Cameron *et al*. from European Genome-phenome Archive (EGA: https://ega-archive.org), under accession number EGA: S00001006136 and EGAS00001006537. The data deposited by Trevino *et al*. and Zhong *et al*. were accessed from Gene Expression Omnibus, under accession number, GEO: GSE162170 and GEO: GSE104276 **(Supplementary Table.13)**.

### Single-Cell RNA Sequencing Data Analysis and Integration

Single-cell RNA sequencing (scRNA-seq) data from six independent studies were processed and integrated using the Seurat package v5.1.0^81^, tidyverse, dplyr in R v4.4.2^82^. Raw count matrices were filtered to include genes expressed in at least 3 cells and cells with a minimum of 200 detected genes. Quality control was performed on each dataset independently, removing cells with >10% mitochondrial gene content and those with total UMIs or detected genes outside the 2.5th to 97.5th percentile range. The quality-controlled datasets were merged and integrated using Seurat’s CCA integration. This involved normalizing each dataset independently using Log-transformation with a scale factor of 10,000, identifying 2,000 variable features per dataset by the variance stabilizing transform (vst) method, and selecting highly variable genes across datasets for integration. Integration anchors were computed and used to create an integrated dataset. Post-integration processing included data scaling with regression of RNA counts, principal component analysis (PCA), and construction of a shared nearest neighbor (SNN) graph using 20 dimensions. Clustering was performed at a resolution of 1.5, resulting in 40 distinct clusters. UMAP was applied for dimensionality reduction and visualization.

### Cell Type Annotation

Cell types were annotated using the scType algorithm^38,39^, which employs predefined lists of positive and negative marker genes for each cell type. This automated approach assigns cell type labels to clusters based on their gene expression profiles. Marker genes used are shown in **Supplementary Table 1**.

### Integration Quality Assessment

To quantitatively assess batch integration and cell type separation, the clustering accuracy with the integration local inverse Simpson’s Index (iLISI) and cell-type LISI (cLISI) scores were calculated using a custom function that employs LISI package^40^. The custom function was validated by cross referencing LISI scores of peripheral blood mononuclear cell (PBMC) data, control (CTRL) and interferon beta stimulated (STIM)^83^, before and after CCA integration described in Stuart and Butler et al., 2018^84^.

### Pseudobulk Differential Gene Expression Analysis

To mitigate the impact of technical noise and uneven cell sampling across donors during differential gene expression (DGE) analysis, a pseudobulk matrix was created by aggregating gene expression data for each cell type within each donor, treating one cell type per donor as a single observation. For temporal DGE analysis, a pseudobulk matrix was created by aggregating gene expression data for each trimester per donor.

Differential gene expression analysis was performed using DESeq2 v1.44.0 on the pseudobulk data^85^. For each cell type/trimester, a binary condition vector was created, labeling the cell type/trimester of interest as the treatment and all others as the control. A DESeqDataSet object was created using the count matrix and sample meta-dataset. Low-count genes (total counts < 10 across all samples) were filtered out. DESeq2 analysis was run using the default parameters. Significant differentially expressed genes (DEGs) were identified by applying stringent thresholds as adjusted p-value < 0.05 and an absolute log2 fold change > 2. Results and data generated from this study were visualized using ComplexHeatmap v2.20.0 and ggplot2 v3.5.1^86^.

### Gene Ontology (GO) term analysis

Over-representation of gene set analysis was conducted by exporting DGE analysis data to Toppgene (https://toppgene.cchmc.org/)^87^. All detected genes for general DGE analysis and 1,027 matrisome genes for matrisome DGE analysis were used as reference background genes for this analysis. The Enrichment Ratio was computed as the proportion of input genes associated with a specific GO term divided by the total number of input genes. The Annotation enrichment was calculated as the percentage of input genes associated with a GO term to the total number of genes associated with that term in the entire annotation database.

### Weighted gene co-expression network analysis

Weighted Gene Co-expression Network Analysis (WGCNA) was employed to identify modules of co-expressed genes and investigate the network properties of our gene of interest (GOI). Using the WGCNA package v1.73^88^, quality control on the expression data was performed, removing genes with excessive missing values or zero variance. The optimal soft-thresholding power was determined by testing a range of powers (1-20) and selecting the lowest power that achieved a scale-free topology fit index (R^2). Using this power value, a signed network was constructed with the blockwiseModules function, setting the minimum module size to 10-30 genes and using a height cut of 0.25 for merging similar modules.

To visualize the network structure around the GOI, the topological overlap matrix (TOM) was calculated using TOMsimilarityFromExpr, and the GOI’s module was subset, and converted to an igraph object^89^. Based on TOM values, the top genes most strongly connected to the GOI were identified. The resulting network was plotted with nodes representing genes and edge widths proportional to connection strengths using custom function.

Pearson correlation coefficient and p-value between the gene of interest (GOI) and co-expressed gene expression levels were calculated using linear regression with ggpubr v0.6.0.

### Pseudotime analysis

Root cells were set as vRG and pseudotime for branching cells was calculated based on their distance from the root cells on the principal graph using monocle3 package v1.3.7 with default parameters^90^.

### ChIP-seq and ATAC-seq analysis

To investigate peaks for histone marks and chromatin accessibility, histone ChIP-seq and ATAC-seq database was accessed from ChIP-Atlas (https://chip-atlas.org/)^62–64^. The combined peak data was visualized using IGV v2.13.1 software.

### Integrated regulatory network analysis

To identify the regulatory network of our gene of interest, the IReNA (Inference of Regulatory Networks using pseudotime-ordered single-cell RNA-seq data) workflow was employed^67^. The motif data was obtained from the TRANSFAC database^91^ and motif-binding transcription factors were extracted. Pseudotime-ordered expression profiles were obtained, filtered for noise, and K-means clustering was performed to group genes with similar expression patterns over pseudotime. Regulatory relationships within grouped genes were identified using Pearson correlation, setting a threshold of |r| > 0.6. To enhance the specificity of our predictions, motifmatchr v1.28.0 was used to detect potential transcription factor (TF) binding sites in the TSS regions of candidate genes. This analysis was performed using the human genome (GRCh38) as a reference. The regulatory relationships of transcription factors and target genes were filtered based on both correlation and motif binding evidence.

### Temporal dynamics of gene expression analysis

To identify genes with significant temporal expression patterns, a custom R function that analyzes gene expression dynamics across developmental stages was developed. The function performs pseudobulk aggregation of single-cell data by donor and timepoint for each cell type, followed by calculation of Pearson’s correlation coefficients and linear regression slopes to quantify expression trends. Genes were filtered for statistical significance (p < 0.05) and strong correlation (|r| > 0.5), with the top genes selected based on their slope magnitudes.

### Cell-to-cell interaction and pathway analysis

To investigate intercellular communication networks, the CellChat package (v1.6.1) was used^55^. A CellChat object was initialized using the matrisome expression matrix and associated meta-data, with cell types as the grouping variable. We utilized the human CellChatDB as the ligand-receptor interaction database. The dataset was subset to include only genes present in CellChatDB. Over-expressed genes and interactions were identified using CellChat’s built-in functions. The communication probability was computed using the trimmed mean approach, and interactions were filtered to include only those involving at least 10 cells. Pathway-level communication probabilities were then calculated.

### Human fetal brain tissue collection

Fetal tissues from elective, normally progressing pregnancies are collected under the Scottish Advanced Fetal Research (SAFeR) study (NCT04613583)^92^. The collection process is approved by one of the 12 Scottish National Health Service Research Ethics Committees (REC 15/NS/0123) and follows the Declaration of Helsinki guidelines. Women seeking elective medical termination of pregnancy were recruited with written informed consent by NHS Grampian research nurses, who operate independently from the research team. There was no alteration in patient treatment or care, and participants could withdraw from the study at any time. The study includes only normally progressing pregnancies, as determined by ultrasound, from women over 16 years of age who speak English and covers gestational ages from 7 to 20 weeks. Grossly abnormal fetuses were excluded, and women experiencing significant distress were not approached. Termination was carried out using RU-486 (Mifepristone) and prostaglandin-induced delivery^93^. Gestational age was confirmed by ultrasound and foot length measurement^94^, and various maternal and fetal data were recorded. For sample collection, fetuses are transported to the laboratory within 30 minutes of delivery, typically intact, and are weighed, measured for crown-rump length (CRL), and sexed by morphology and PCR confirmation of the Y chromosome. Collected brain tissues were fixed in 4% PFA in phosphate-buffered overnight at 4⁰C followed by preservation in 15% sucrose in PBS at 4⁰C overnight followed by 30% sucrose-sodium azide solution (7.7mM NaN_3_). Nine samples were used in this study, include GW9 (Male), GW11 (Male), GW12 (Female), GW14 (Male), GW15 (female), two GW16 (both female) and two GW17 (both male).

### Immunofluorescence staining

The fixed prefrontal cortical samples were embedded in OCT compound and frozen at -80⁰C, prior to being cryosectioned at a 20 µm thickness (Leica 1850 croystats). Sections were retrieved onto Superfrost glass slides and stored at -20⁰C. Sections were washed in tris-buffered saline (TBS, 137 mM NaCl, 25mM Tris-HCl and 2.7mM KCl), and antigen retrieval was performed using Target retrieval solution (Dako, S1699). Sections were permeabilized with 0.05% Triton-X-100 in TBS for 15 minutes at RT and blocked with blocking buffer at room temperature for 1 hour. Primary antibodies were diluted in blocking buffer and incubated with sections over two nights at 4⁰C. Sections were washed three times with 0.05% Triton X-100 in TBS and incubated with secondary antibodies diluted in blocking buffer at 4⁰C for 2 hours.

Sections were washed three times and allowed to dry before treating with Vector TrueView Autofluorescence Quenching Kit with DAPI (Vector, SP-8500), and coverslips were mounted. Antibodies used in this study are listed in **Supplementary Table 14**.

### Imaging and Image analysis

A Zeiss LSM880 confocal microscope and Airyscan Fast with 20X objective was used to acquire images. Z-stacks were collected at 1µm intervals and image tiles were automatically processed by inbuilt Zeiss ZEN 3.0 software. Object-based colocalization analysis and semi-supervised cell count were performed using Colocalization Image Creator and Colocalization Cell Counter plugins on ImageJ^95^. Total number of DAPI^+^ cells were determined using automatic object counter (radius at 5 micron and noise tolerance at 70) followed by visual inspection of cell counts. Number of HOPX^+^, S100β^+^ and OLIG2^+^ cells was determined by generating colocalization binary image with DAPI. For Galectin-3 cell count, Galectin-3 grayscale image was merged with DAPI^+^ binary image and DAPI and HOPX colocalization binary image each. Using this, number of Galectin-3^+^ cells and Galectin-3^+^ and HOPX^+^ cells was identified. The statistical analysis of mean count differences was performed on three biological replicates for HOPX/Galectin-3 staining and four biological replicates for OLIG2 and S100β staining across GW 15-17.

### Code availability

All codes used in this study are available at https://github.com/KANGBERGLAB/matrisome-meta-analysis/tree/main

## Acknowledgments

We thank members of the Kang and Berg laboratories for comments and suggestions. We thank the members of Fowler Lab for support in fetal sample acquisition. This work was supported by grants from The Humane Research Trust Les Rhoades PhD scholarship (to E.K.), The Academy of Medical Sciences Springboard (SBF007\100169 to E.K.) and Biological Sciences Research Council (BB/W008068/1 to D.A.B.). The SAFeR study was funded by the UK Medical Research Council (MR/L010011/1 and MR/P011535/1) and the EU’s Horizon 2020 research and innovation programme under the Marie Skłodowska-Curie project PROTECTED (grant agreement number 722634) and FREIA and INITIALISE projects (grant agreement numbers 825100 and 101094099 respectively) to PAF.

## Author Contributions

E.K., D.A.B. and D.H.G. conceived the project and wrote the paper. D.H.G. led and contributed to all aspects of the study. M.Z.K.A. and M.M. provided guidance and suggestions on data analysis. P.A.F. contributed to obtaining fetal brain samples. All authors commented on and approved publication of manuscript.

## References

1 Jovanov Milosevic, N., Judas, M., Aronica, E. & Kostovic, I. Neural ECM in laminar organization and connectivity development in healthy and diseased human brain. Prog Brain Res 214, 159–178 (2014). 10.1016/B978-0-444-63486-3.00007-4

2 Long, K. R. & Huttner, W. B. The Role of the Extracellular Matrix in Neural Progenitor Cell Proliferation and Cortical Folding During Human Neocortex Development. Front Cell Neurosci 15, 804649 (2021). 10.3389/fncel.2021.804649

3 Sykova, E. & Nicholson, C. Diffusion in brain extracellular space. Physiol Rev 88, 1277–1340 (2008). 10.1152/physrev.00027.2007

4 Fietz, S. A. et al. OSVZ progenitors of human and ferret neocortex are epithelial-like and expand by integrin signaling. Nat Neurosci 13, 690–699 (2010). 10.1038/nn.2553

5 Long, K., Moss, L., Laursen, L., Boulter, L. & Ffrench-Constant, C. Integrin signalling regulates the expansion of neuroepithelial progenitors and neurogenesis via Wnt7a and Decorin. Nat Commun 7, 10354 (2016). 10.1038/ncomms10354

6 Loulier, K. et al. beta1 integrin maintains integrity of the embryonic neocortical stem cell niche. PLoS Biol 7, e1000176 (2009). 10.1371/journal.pbio.1000176

7 Radakovits, R., Barros, C. S., Belvindrah, R., Patton, B. & Muller, U. Regulation of radial glial survival by signals from the meninges. J Neurosci 29, 7694–7705 (2009). 10.1523/JNEUROSCI.5537-08.2009

8 Chen, Z. L., Haegeli, V., Yu, H. & Strickland, S. Cortical deficiency of laminin gamma1 impairs the AKT/GSK-3beta signaling pathway and leads to defects in neurite outgrowth and neuronal migration. Dev Biol 327, 158–168 (2009). 10.1016/j.ydbio.2008.12.006

9 Lander, A. D., Fujii, D. K. & Reichardt, L. F. Purification of a factor that promotes neurite outgrowth: isolation of laminin and associated molecules. J Cell Biol 101, 898–913 (1985). 10.1083/jcb.101.3.898

10 Ma, W. et al. Cell-extracellular matrix interactions regulate neural differentiation of human embryonic stem cells. BMC Dev Biol 8, 90 (2008). 10.1186/1471-213X-8-90

11 Myers, J. P., Santiago-Medina, M. & Gomez, T. M. Regulation of axonal outgrowth and pathfinding by integrin-ECM interactions. Dev Neurobiol 71, 901–923 (2011). 10.1002/dneu.20931

12 Long, K. R. et al. Extracellular Matrix Components HAPLN1, Lumican, and Collagen I Cause Hyaluronic Acid-Dependent Folding of the Developing Human Neocortex. Neuron 99, 702–719 e706 (2018). 10.1016/j.neuron.2018.07.013

13 Hong, S. E. et al. Autosomal recessive lissencephaly with cerebellar hypoplasia is associated with human RELN mutations. Nat Genet 26, 93–96 (2000). 10.1038/79246

14 Eriksson, S. H. et al. Persistent reelin-expressing Cajal-Retzius cells in polymicrogyria. Brain 124, 1350–1361 (2001). 10.1093/brain/124.7.1350

15 Bosiacki, M. et al. Perineuronal Nets and Their Role in Synaptic Homeostasis. Int J Mol Sci 20 (2019). 10.3390/ijms20174108

16 Dityatev, A., Schachner, M. & Sonderegger, P. The dual role of the extracellular matrix in synaptic plasticity and homeostasis. Nat Rev Neurosci 11, 735–746 (2010). 10.1038/nrn2898

17 Matsuda, K. et al. Cbln1 is a ligand for an orphan glutamate receptor delta2, a bidirectional synapse organizer. Science 328, 363–368 (2010). 10.1126/science.1185152

18 Hynes, R. O. & Naba, A. Overview of the matrisome--an inventory of extracellular matrix constituents and functions. Cold Spring Harb Perspect Biol 4, a004903 (2012). 10.1101/cshperspect.a004903

19 D’Arcangelo, G. et al. A protein related to extracellular matrix proteins deleted in the mouse mutant reeler. Nature 374, 719–723 (1995). 10.1038/374719a0

20 Devisme, L. et al. Cobblestone lissencephaly: neuropathological subtypes and correlations with genes of dystroglycanopathies. Brain 135, 469–482 (2012). 10.1093/brain/awr357

21 Radner, S. et al. beta2 and gamma3 laminins are critical cortical basement membrane components: ablation of Lamb2 and Lamc3 genes disrupts cortical lamination and produces dysplasia. Dev Neurobiol 73, 209–229 (2013). 10.1002/dneu.22057

22 Barak, T. et al. Recessive LAMC3 mutations cause malformations of occipital cortical development. Nat Genet 43, 590–594 (2011). 10.1038/ng.836

23 Hubert, T., Grimal, S., Carroll, P. & Fichard-Carroll, A. Collagens in the developing and diseased nervous system. Cell Mol Life Sci 66, 1223–1238 (2009). 10.1007/s00018-008-8561-9

24 Amin, S. & Borrell, V. The Extracellular Matrix in the Evolution of Cortical Development and Folding. Front Cell Dev Biol 8, 604448 (2020). 10.3389/fcell.2020.604448

25 Fietz, S. A. et al. Transcriptomes of germinal zones of human and mouse fetal neocortex suggest a role of extracellular matrix in progenitor self-renewal. Proc Natl Acad Sci U S A 109, 11836–11841 (2012). 10.1073/pnas.1209647109

26 Vinsland, E. & Linnarsson, S. Single-cell RNA-sequencing of mammalian brain development: insights and future directions. Development 149 (2022). 10.1242/dev.200180

27 Sonrel, A. et al. Meta-analysis of (single-cell method) benchmarks reveals the need for extensibility and interoperability. Genome Biol 24, 119 (2023). 10.1186/s13059-023-02962-5

28 Zhong, S. et al. A single-cell RNA-seq survey of the developmental landscape of the human prefrontal cortex. Nature 555, 524–528 (2018). 10.1038/nature25980

29 Trevino, A. E. et al. Chromatin and gene-regulatory dynamics of the developing human cerebral cortex at single-cell resolution. Cell 184, 5053–5069 e5023 (2021). 10.1016/j.cell.2021.07.039

30 Cameron, D. et al. Single-Nuclei RNA Sequencing of 5 Regions of the Human Prenatal Brain Implicates Developing Neuron Populations in Genetic Risk for Schizophrenia. Biol Psychiatry 93, 157–166 (2023). 10.1016/j.biopsych.2022.06.033

31 van Bruggen, D. et al. Developmental landscape of human forebrain at a single-cell level identifies early waves of oligodendrogenesis. Dev Cell 57, 1421–1436 e1425 (2022). 10.1016/j.devcel.2022.04.016

32 Bhaduri, A. et al. An atlas of cortical arealization identifies dynamic molecular signatures. Nature 598, 200–204 (2021). 10.1038/s41586-021-03910-8

33 Eze, U. C., Bhaduri, A., Haeussler, M., Nowakowski, T. J. & Kriegstein, A. R. Single-cell atlas of early human brain development highlights heterogeneity of human neuroepithelial cells and early radial glia. Nat Neurosci 24, 584–594 (2021). 10.1038/s41593-020-00794-1

34 Shao, X. et al. MatrisomeDB 2.0: 2023 updates to the ECM-protein knowledge database. Nucleic Acids Res 51, D1519–D1530 (2023). 10.1093/nar/gkac1009

35 Banerjee-Basu, S. & Packer, A. SFARI Gene: an evolving database for the autism research community. Dis Model Mech 3, 133–135 (2010). 10.1242/dmm.005439

36 Gonzalez-Mantilla, A. J., Moreno-De-Luca, A., Ledbetter, D. H. & Martin, C. L. A Cross-Disorder Method to Identify Novel Candidate Genes for Developmental Brain Disorders. JAMA Psychiatry 73, 275–283 (2016). 10.1001/jamapsychiatry.2015.2692

37 Kochinke, K. et al. Systematic Phenomics Analysis Deconvolutes Genes Mutated in Intellectual Disability into Biologically Coherent Modules. Am J Hum Genet 98, 149–164 (2016). 10.1016/j.ajhg.2015.11.024

38 Ianevski, A., Giri, A. K. & Aittokallio, T. Fully-automated and ultra-fast cell-type identification using specific marker combinations from single-cell transcriptomic data. Nat Commun 13, 1246 (2022). 10.1038/s41467-022-28803-w

39 Oh, S. et al. HGNChelper: identification and correction of invalid gene symbols for human and mouse. F1000Res 9, 1493 (2020). 10.12688/f1000research.28033.2

40 Korsunsky, I. et al. Fast, sensitive and accurate integration of single-cell data with Harmony. Nat Methods 16, 1289–1296 (2019). 10.1038/s41592-019-0619-0

41 Walker, F. R. et al. Dynamic structural remodelling of microglia in health and disease: a review of the models, the signals and the mechanisms. Brain Behav Immun 37, 1–14 (2014). 10.1016/j.bbi.2013.12.010

42 Faissner, A. & Reinhard, J. The extracellular matrix compartment of neural stem and glial progenitor cells. Glia 63, 1330–1349 (2015). 10.1002/glia.22839

43 Hirota, Y. & Nakajima, K. Control of Neuronal Migration and Aggregation by Reelin Signaling in the Developing Cerebral Cortex. Front Cell Dev Biol 5, 40 (2017). 10.3389/fcell.2017.00040

44 Bonnans, C., Chou, J. & Werb, Z. Remodelling the extracellular matrix in development and disease. Nat Rev Mol Cell Biol 15, 786–801 (2014). 10.1038/nrm3904

45 Gao, Q., Mok, H. P. & Zhuang, J. Secreted modular calcium-binding proteins in pathophysiological processes and embryonic development. Chin Med J (Engl) 132, 2476–2484 (2019). 10.1097/CM9.0000000000000472

46 Gan, K. J. & Sudhof, T. C. SPARCL1 Promotes Excitatory But Not Inhibitory Synapse Formation and Function Independent of Neurexins and Neuroligins. J Neurosci 40, 8088–8102 (2020). 10.1523/JNEUROSCI.0454-20.2020

47 Niu, X., Zhang, F., Gu, W., Zhang, B. & Chen, X. FBLN2 is associated with Goldenhar syndrome and is essential for cranial neural crest cell development. Ann N Y Acad Sci 1537, 113–128 (2024). 10.1111/nyas.15183

48 Fels, E. et al. Role of LGI1 protein in synaptic transmission: From physiology to pathology. Neurobiol Dis 160, 105537 (2021). 10.1016/j.nbd.2021.105537

49 Assimacopoulos, S., Grove, E. A. & Ragsdale, C. W. Identification of a Pax6-dependent epidermal growth factor family signaling source at the lateral edge of the embryonic cerebral cortex. J Neurosci 23, 6399–6403 (2003). 10.1523/JNEUROSCI.23-16-06399.2003

50 Taetzsch, T., Brayman, V. L. & Valdez, G. FGF binding proteins (FGFBPs): Modulators of FGF signaling in the developing, adult, and stressed nervous system. Biochim Biophys Acta Mol Basis Dis 1864, 2983–2991 (2018). 10.1016/j.bbadis.2018.06.009

51 Zhang, X. F. et al. The function of the inter-alpha-trypsin inhibitors in the development of disease. Front Med (Lausanne) 11, 1432224 (2024). 10.3389/fmed.2024.1432224

52 Rapacioli, M., Palma, V. & Flores, V. Morphogenetic and Histogenetic Roles of the Temporal-Spatial Organization of Cell Proliferation in the Vertebrate Corticogenesis as Revealed by Inter-specific Analyses of the Optic Tectum Cortex Development. Front Cell Neurosci 10, 67 (2016). 10.3389/fncel.2016.00067

53 Abdulmalek, S. et al. Midkine is upregulated in the hippocampus following both spatial and olfactory reward association learning and enhances memory. J Neurochem 168, 2832–2847 (2024). 10.1111/jnc.16151

54 Zheng, Y. Z. & Liang, L. High expression of PXDN is associated with poor prognosis and promotes proliferation, invasion as well as migration in ovarian cancer. Ann Diagn Pathol 34, 161–165 (2018). 10.1016/j.anndiagpath.2018.03.002

55 Jin, S. et al. Inference and analysis of cell-cell communication using CellChat. Nat Commun 12, 1088 (2021). 10.1038/s41467-021-21246-9

56 Pufe, T., Bartscher, M., Petersen, W., Tillmann, B. & Mentlein, R. Expression of pleiotrophin, an embryonic growth and differentiation factor, in rheumatoid arthritis. Arthritis Rheum 48, 660–667 (2003). 10.1002/art.10839

57 Herradon, G. & Perez-Garcia, C. Targeting midkine and pleiotrophin signalling pathways in addiction and neurodegenerative disorders: recent progress and perspectives. Br J Pharmacol 171, 837–848 (2014). 10.1111/bph.12312

58 Yildirim, B., Kulak, K. & Bilir, A. Midkine: A Cancer Biomarker Candidate and Innovative Therapeutic Approaches. Eur J Breast Health 20, 167–177 (2024). 10.4274/ejbh.galenos.2024.2024-4-7

59 Perez-Branguli, F. et al. Reverse Signaling by Semaphorin-6A Regulates Cellular Aggregation and Neuronal Morphology. PLoS One 11, e0158686 (2016). 10.1371/journal.pone.0158686

60 Kerjan, G. et al. The transmembrane semaphorin Sema6A controls cerebellar granule cell migration. Nat Neurosci 8, 1516–1524 (2005). 10.1038/nn1555

61 Pollen, A. A. et al. Molecular identity of human outer radial glia during cortical development. Cell 163, 55–67 (2015). 10.1016/j.cell.2015.09.004

62 Zou, Z., Ohta, T. & Oki, S. ChIP-Atlas 3.0: a data-mining suite to explore chromosome architecture together with large-scale regulome data. Nucleic Acids Res 52, W45–W53 (2024). 10.1093/nar/gkae358

63 Zou, Z., Ohta, T., Miura, F. & Oki, S. ChIP-Atlas 2021 update: a data-mining suite for exploring epigenomic landscapes by fully integrating ChIP-seq, ATAC-seq and Bisulfite-seq data. Nucleic Acids Res 50, W175–W182 (2022). 10.1093/nar/gkac199

64 Oki, S. et al. ChIP-Atlas: a data-mining suite powered by full integration of public ChIP-seq data. EMBO Rep 19 (2018). 10.15252/embr.201846255

65 Meijer, M. et al. Epigenomic priming of immune genes implicates oligodendroglia in multiple sclerosis susceptibility. Neuron 110, 1193–1210 e1113 (2022). 10.1016/j.neuron.2021.12.034

66 Yu, Y. et al. H3K27me3-H3K4me1 transition at bivalent promoters instructs lineage specification in development. Cell Biosci 13, 66 (2023). 10.1186/s13578-023-01017-3

67 Jiang, J. et al. IReNA: Integrated regulatory network analysis of single-cell transcriptomes and chromatin accessibility profiles. iScience 25, 105359 (2022). 10.1016/j.isci.2022.105359

68 Deloulme, J. C. et al. Nuclear expression of S100B in oligodendrocyte progenitor cells correlates with differentiation toward the oligodendroglial lineage and modulates oligodendrocytes maturation. Mol Cell Neurosci 27, 453–465 (2004). 10.1016/j.mcn.2004.07.008

69 Hernandez-Ortega, K. et al. S100B actions on glial and neuronal cells in the developing brain: an overview. Front Neurosci 18, 1425525 (2024). 10.3389/fnins.2024.1425525

70 Lawrence, A. R. et al. Microglia maintain structural integrity during fetal brain morphogenesis. Cell 187, 962–980 e919 (2024). 10.1016/j.cell.2024.01.012

71 Crouch, E. E. et al. Ensembles of endothelial and mural cells promote angiogenesis in prenatal human brain. Cell 185, 3753–3769 e3718 (2022). 10.1016/j.cell.2022.09.004

72 Zhang, D. et al. Neuregulin-3 (NRG3): a novel neural tissue-enriched protein that binds and activates ErbB4. Proc Natl Acad Sci U S A 94, 9562–9567 (1997). 10.1073/pnas.94.18.9562

73 Muller, T. et al. Neuregulin 3 promotes excitatory synapse formation on hippocampal interneurons. EMBO J 37 (2018). 10.15252/embj.201798858

74 Long, K. R. & Huttner, W. B. How the extracellular matrix shapes neural development. Open Biol 9, 180216 (2019). 10.1098/rsob.180216

75 Ferent, J., Zaidi, D. & Francis, F. Extracellular Control of Radial Glia Proliferation and Scaffolding During Cortical Development and Pathology. Front Cell Dev Biol 8, 578341 (2020). 10.3389/fcell.2020.578341

76 Wang, L. et al. Molecular and cellular dynamics of the developing human neocortex at single-cell resolution. bioRxiv (2024). 10.1101/2024.01.16.575956

77 Assir, M. Z. K. et al. IL-17A Alters Human Cortical Development in a 3D Ex Vivo Model of Maternal Immune Activation. bioRxiv, 2024.2010.2030.621011 (2024). 10.1101/2024.10.30.621011

78 Sarkar, A. & Wintrode, P. L. Effects of glycosylation on the stability and flexibility of a metastable protein: the human serpin alpha(1)-antitrypsin. Int J Mass Spectrom 302, 69–75 (2011). 10.1016/j.ijms.2010.08.003

79 Kellman, B. P. & Lewis, N. E. Big-Data Glycomics: Tools to Connect Glycan Biosynthesis to Extracellular Communication. Trends Biochem Sci 46, 284–300 (2021). 10.1016/j.tibs.2020.10.004

80 Naba, A. et al. The matrisome: in silico definition and in vivo characterization by proteomics of normal and tumor extracellular matrices. Mol Cell Proteomics 11, M111 014647 (2012). 10.1074/mcp.M111.014647

81 Hao, Y. et al. Integrated analysis of multimodal single-cell data. Cell 184, 3573–3587 e3529 (2021). 10.1016/j.cell.2021.04.048

82 Hao, Y. et al. Dictionary learning for integrative, multimodal and scalable single-cell analysis. Nat Biotechnol 42, 293–304 (2024). 10.1038/s41587-023-01767-y

83 Kang, H. M. et al. Multiplexed droplet single-cell RNA-sequencing using natural genetic variation. Nat Biotechnol 36, 89–94 (2018). 10.1038/nbt.4042

84 Stuart, T. et al. Comprehensive Integration of Single-Cell Data. Cell 177, 1888–1902 e1821 (2019). 10.1016/j.cell.2019.05.031

85 Love, M. I., Huber, W. & Anders, S. Moderated estimation of fold change and dispersion for RNA-seq data with DESeq2. Genome Biol 15, 550 (2014). 10.1186/s13059-014-0550-8

86 Wickham, H. in Use R!, 1 online resource (XVI, 260 pages 232 illustrations, 140 illustrations in color (Springer International Publishing : Imprint: Springer,, Cham, 2016).

87 Chen, J., Bardes, E. E., Aronow, B. J. & Jegga, A. G. ToppGene Suite for gene list enrichment analysis and candidate gene prioritization. Nucleic Acids Res 37, W305–311 (2009). 10.1093/nar/gkp427

88 Langfelder, P. & Horvath, S. WGCNA: an R package for weighted correlation network analysis. BMC Bioinformatics 9, 559 (2008). 10.1186/1471-2105-9-559

89 Zhang, B. & Horvath, S. A general framework for weighted gene co-expression network analysis. Stat Appl Genet Mol Biol 4, Article17 (2005). 10.2202/1544-6115.1128

90 Trapnell, C. et al. The dynamics and regulators of cell fate decisions are revealed by pseudotemporal ordering of single cells. Nat Biotechnol 32, 381–386 (2014). 10.1038/nbt.2859

91 Matys, V. et al. TRANSFAC and its module TRANSCompel: transcriptional gene regulation in eukaryotes. Nucleic Acids Res 34, D108–110 (2006). 10.1093/nar/gkj143

92 Bongaerts, E. et al. Maternal exposure to ambient black carbon particles and their presence in maternal and fetal circulation and organs: an analysis of two independent population-based observational studies. Lancet Planet Health 6, e804–e811 (2022). 10.1016/S2542-5196(22)00200-5

93 O’Shaughnessy, P. J. et al. Developmental changes in human fetal testicular cell numbers and messenger ribonucleic acid levels during the second trimester. J Clin Endocrinol Metab 92, 4792–4801 (2007). 10.1210/jc.2007-1690

94 Evtouchenko, L., Studer, L., Spenger, C., Dreher, E. & Seiler, R. W. A mathematical model for the estimation of human embryonic and fetal age. Cell Transplant 5, 453–464 (1996). 10.1177/096368979600500404

95 Lunde, A. & Glover, J. C. A versatile toolbox for semi-automatic cell-by-cell object-based colocalization analysis. Sci Rep 10, 19027 (2020). 10.1038/s41598-020-75835-7

