## supplementary information for "Deciphering Cell-Type and Temporal-Specific Matrisome Expression Signatures in Human Cortical Development and Neurodevelopmental Disorders via scRNA-Seq Meta-Analysis"

**The PDF file includes:**

Supplementary Figures 1 to 6

**Other Supplementary Information for this manuscript includes the following:**

Supplementary Tables 1 to 14

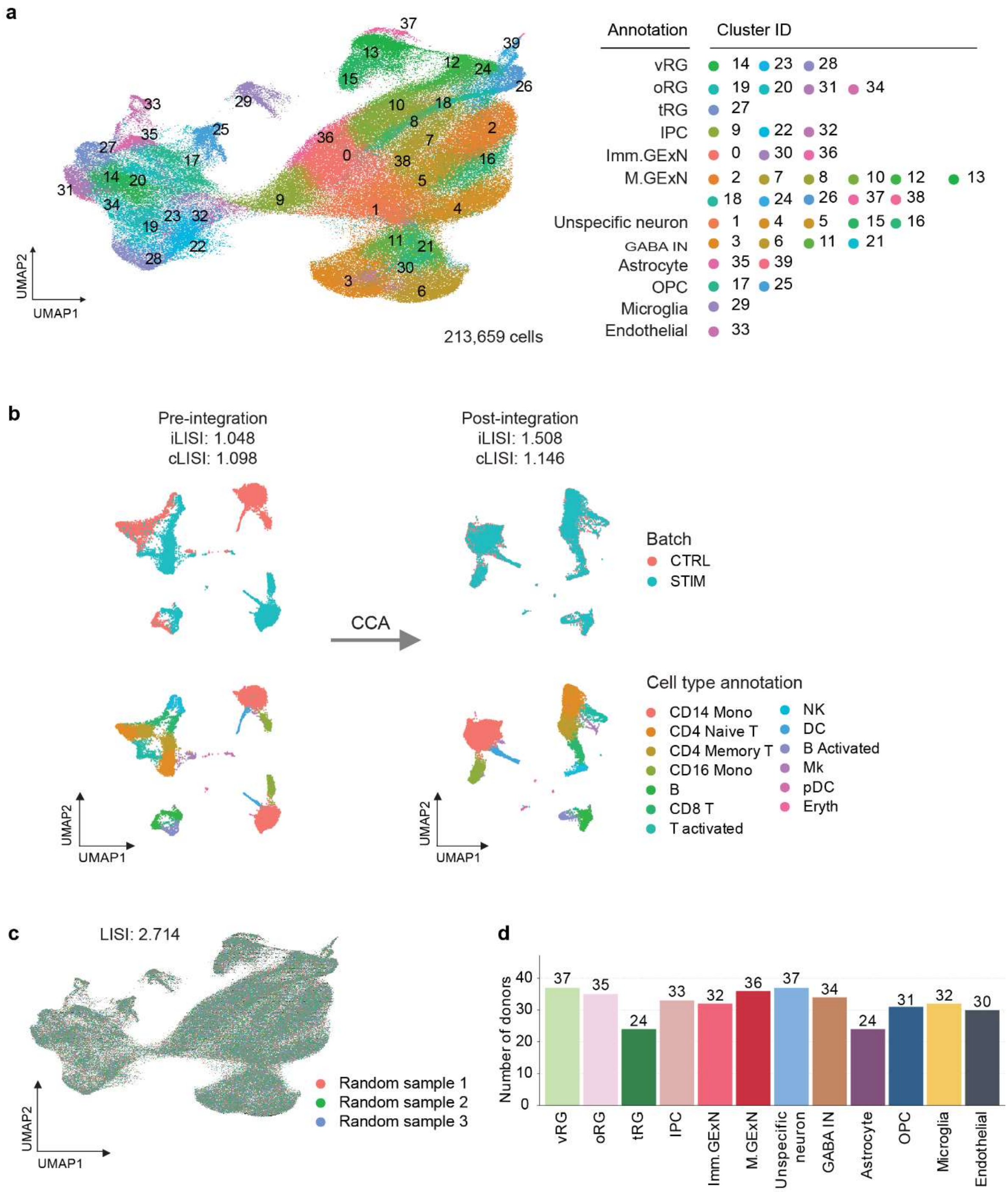


**Supplementary Fig. 1: Cell type annotation of meta-data and validation of integration**

**a** Unsupervised cell type annotation: 40 clusters were identified in the meta-data, and each cluster was annotated based on cell type using the scType pipeline (left). A table presents the corresponding cell type for each cluster ID (right).

**b** UMAP visualization of two groups of peripheral blood mononuclear cell (PBMC) data—control (CTRL) and interferon beta-stimulated (STIM)—before and after CCA integration. The Integration Local Inverse Simpson's Index (iLISI) and Cell-Type LISI (cLISI) scores are displayed before and after integration. Notably, iLISI increased from 1.048 to 1.508 following integration.

**c** UMAP visualization of the scRNA-seq meta-data with random sampling annotations (Sample 1–3) representing an ideal batch integration. The Local Inverse Simpson's Index (LISI) for perfect batch mixing is shown as the maximum score reference for this dataset.

**d** Bar graph displaying the number of donors per cell type.

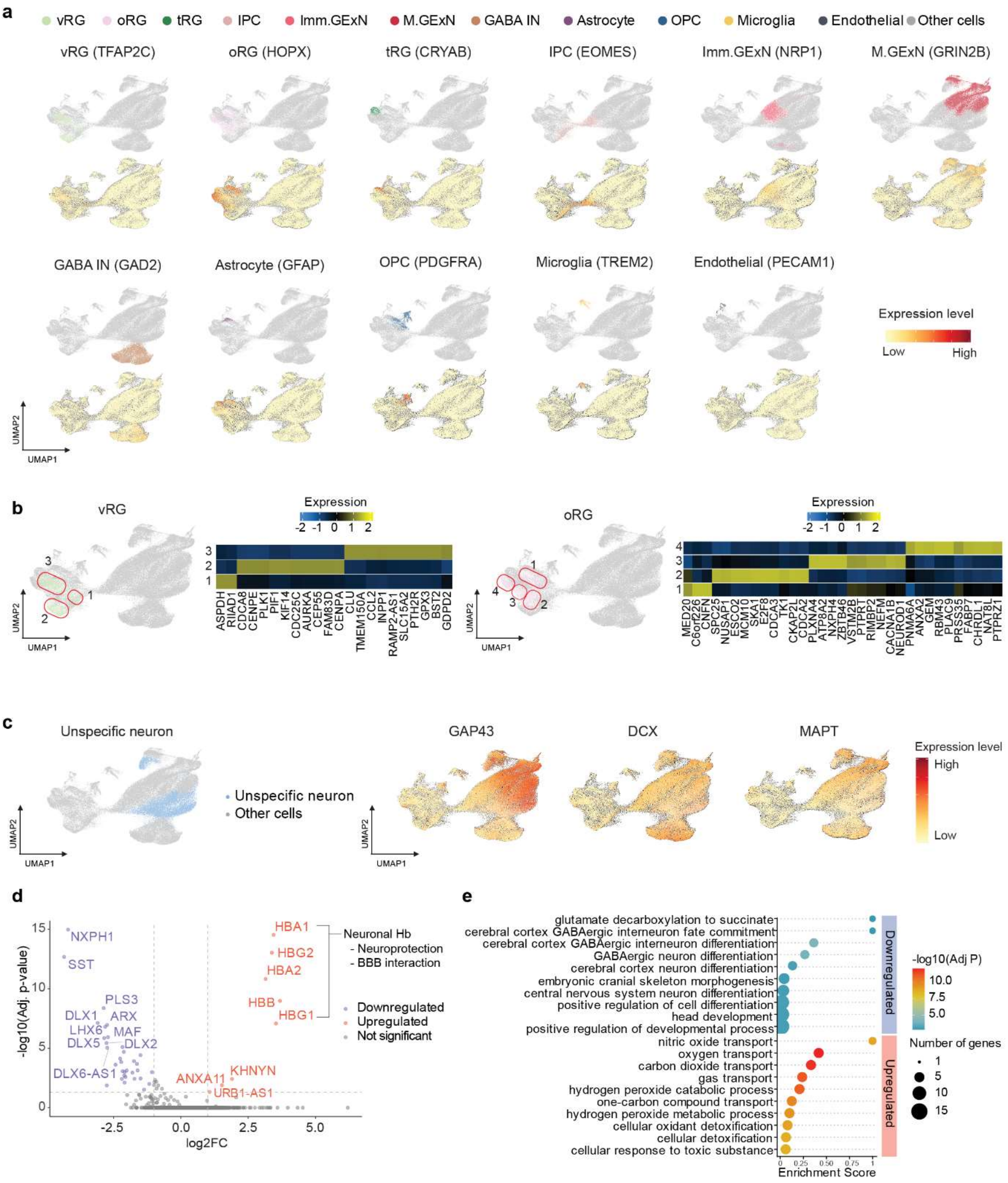


### Supplementary Figure 2: Validation and Characterization of Cell Type Annotation

**a** UMAP visualization of the scRNA-seq meta-data highlighting all cell type annotations and known cell type marker genes. Expression levels of marker genes are shown on a scaled color gradient.

**b** UMAP visualization of the scRNA-seq meta-data highlighting distinct vRG clusters (left) and oRG clusters (right). Red circles indicate each cluster of cells, labeled with cluster numbers. A heatmap displays differentially expressed genes between the clusters. The heatmap represents DEG expression levels across distinct clusters, where each column corresponds to a gene, and each row represents a cell type ($log_2$ fold-change > 1, adjusted p-value < 0.05).

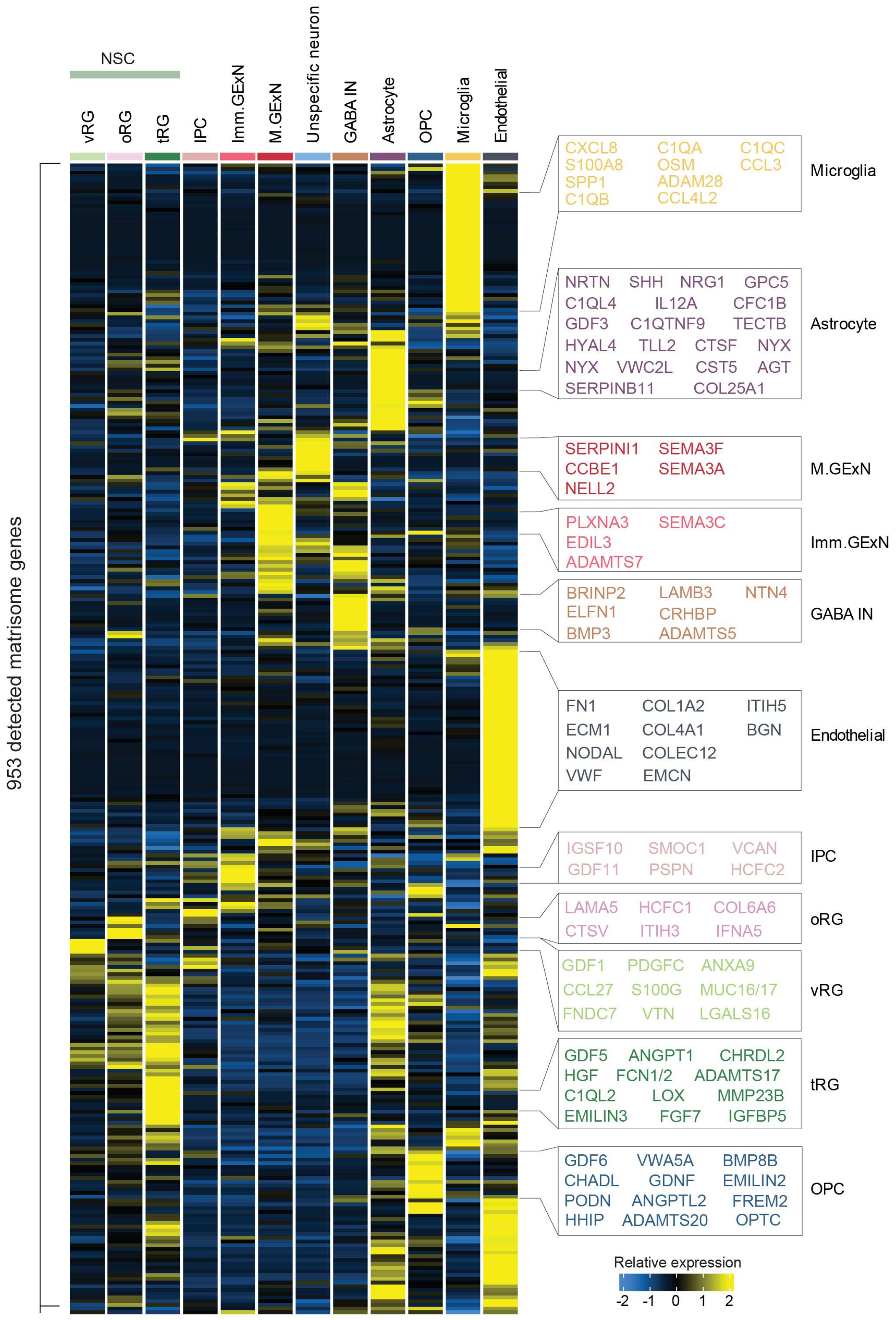


**Supplementary Fig. 3: Matrisome expression across the various cell types**

Heatmap of relative expression levels of 953 matrisome genes across 12 cell types. Each row represents a gene, and each column corresponds to a cell type, highlighting cell type-specific signatures of matrisome gene.

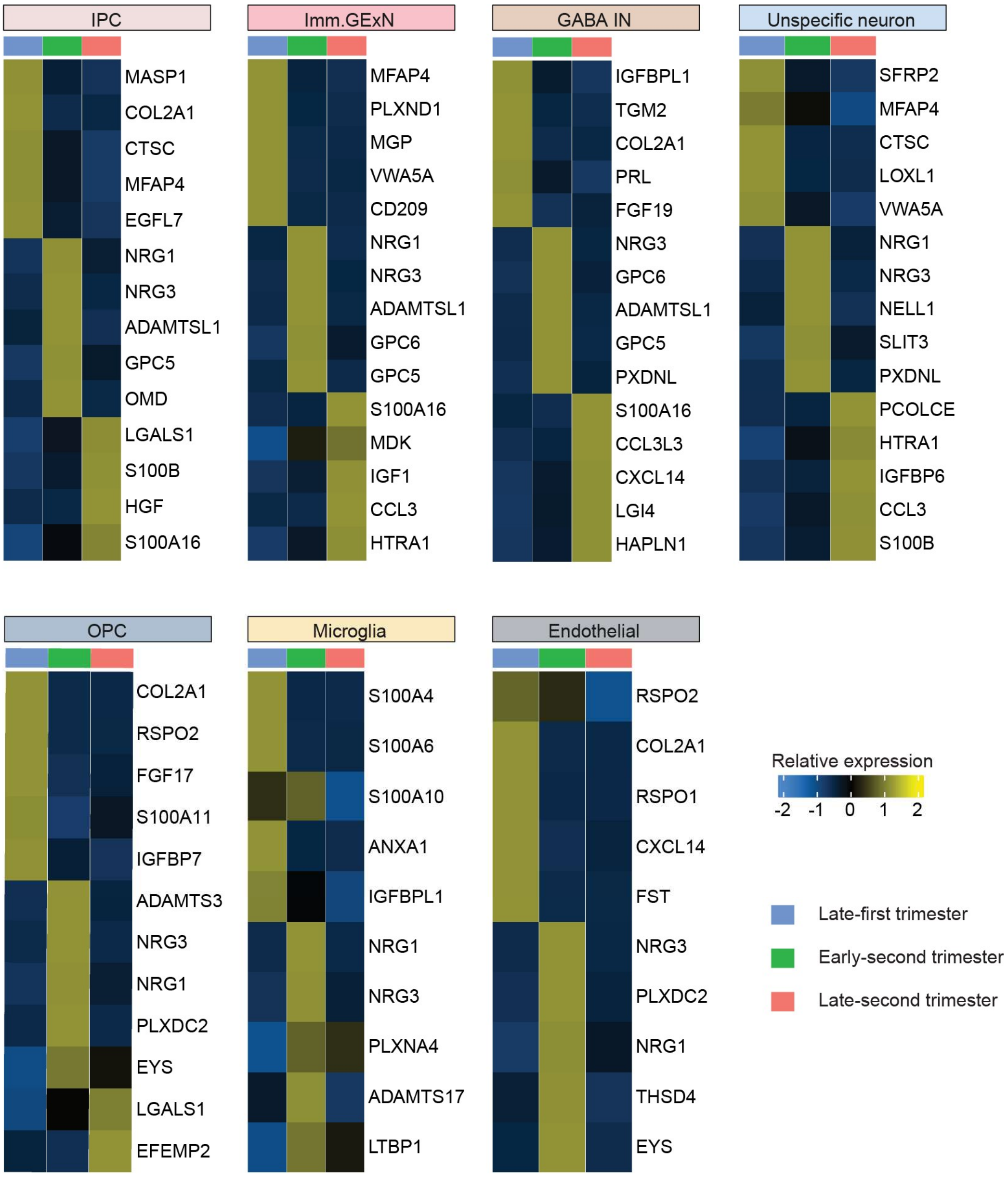


**Supplementary Fig. 4: Temporal-specific matrisome signatures in each cell type**

Heatmap displaying the expression levels of the top matrisome genes across the late first trimester, early second trimester, and mid-second trimester in different cell types. Each row represents a gene, and each column corresponds to a developmental period, highlighting cell type-specific marker genes (ranked by $\log_2$ fold-change, adjusted p-value < 0.05).

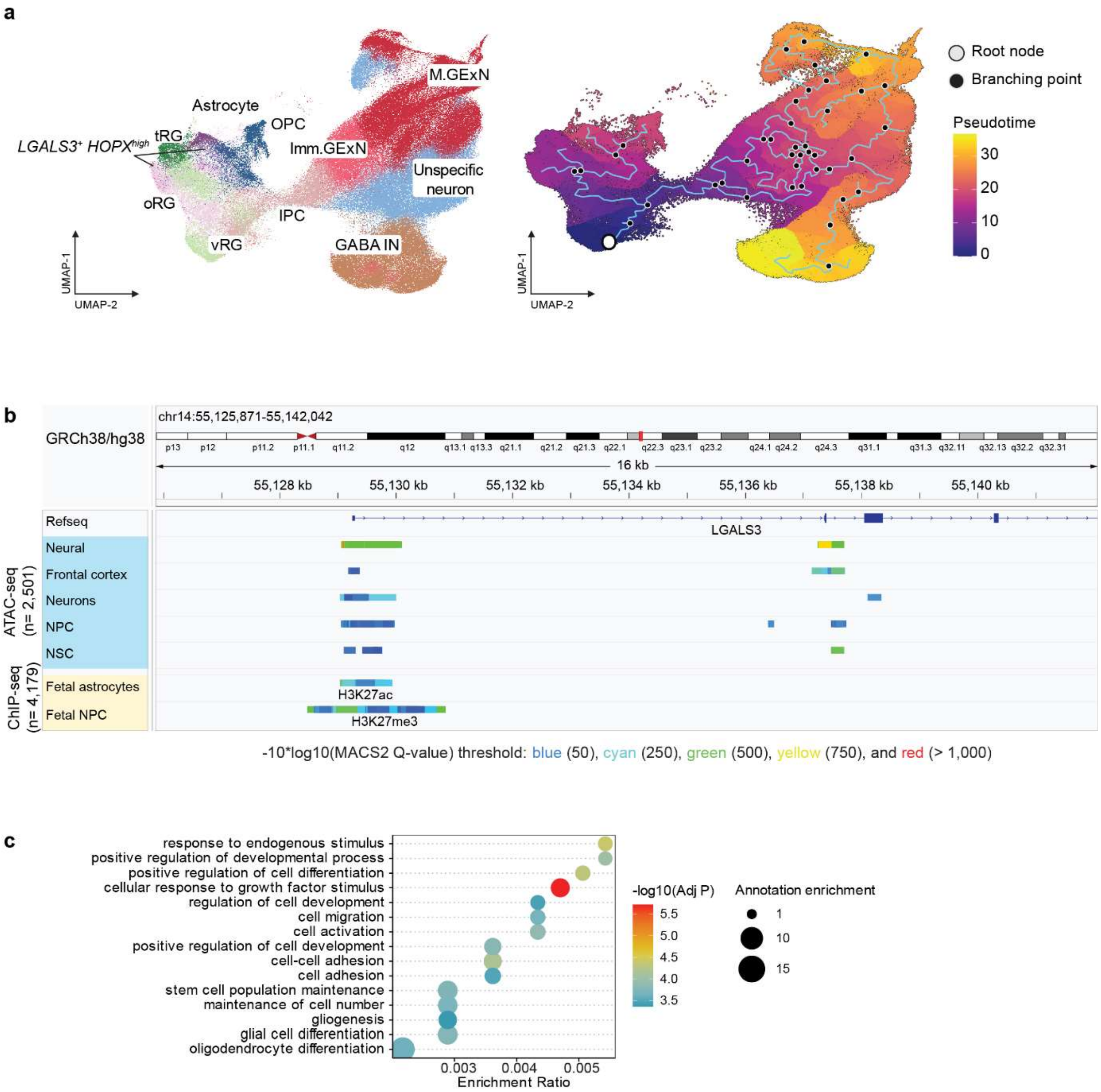


**Supplementary Fig. 5: Characterization of LGALS3$^+$ HOPX$^{high}$ cells and genomic regulatory landscape of LGALS3**

**a** UMAP visualizations of scRNA-seq meta-data in the developing cortex. Left: UMAP showing cell type annotations and highlighting and LGALS3$^+$ HOPX$^{high}$ cells. Right: UMAP illustrating the developmental trajectory inferred using pseudotime analysis. Pseudotime, computed from gene expression profiles of each cluster in the meta-data, is represented by a color scale. The root node (vRG) and key branching points of the developmental trajectory are indicated.

**b** Chromatin accessibility and histone modification profiles at the LGALS3 locus. Genomic tracks display chromatin accessibility (ATAC-seq) and histone modifications (H3K27ac, H3K27me3) across different cell types. The tracks represent signal peaks from 2,501 ATAC-seq datasets and 4,179 ChIP-seq datasets at chr14:55,125,871-55,142,042 (16 kb). Signal strength is represented by statistical significance and color intensity based on MACS2 Q-values (-10*log10 transformed), with thresholds indicated by blue (50), cyan (250), green (500), yellow (750), and red (>1,000).

**c** Gene Ontology (GO) term analysis of Group 1 transcription activators regulating LGALS3. The x-axis represents the enrichment score, dot size indicates the number of genes in each annotation, and the $-\log_{10}$ adjusted p-value is represented by the color scale.

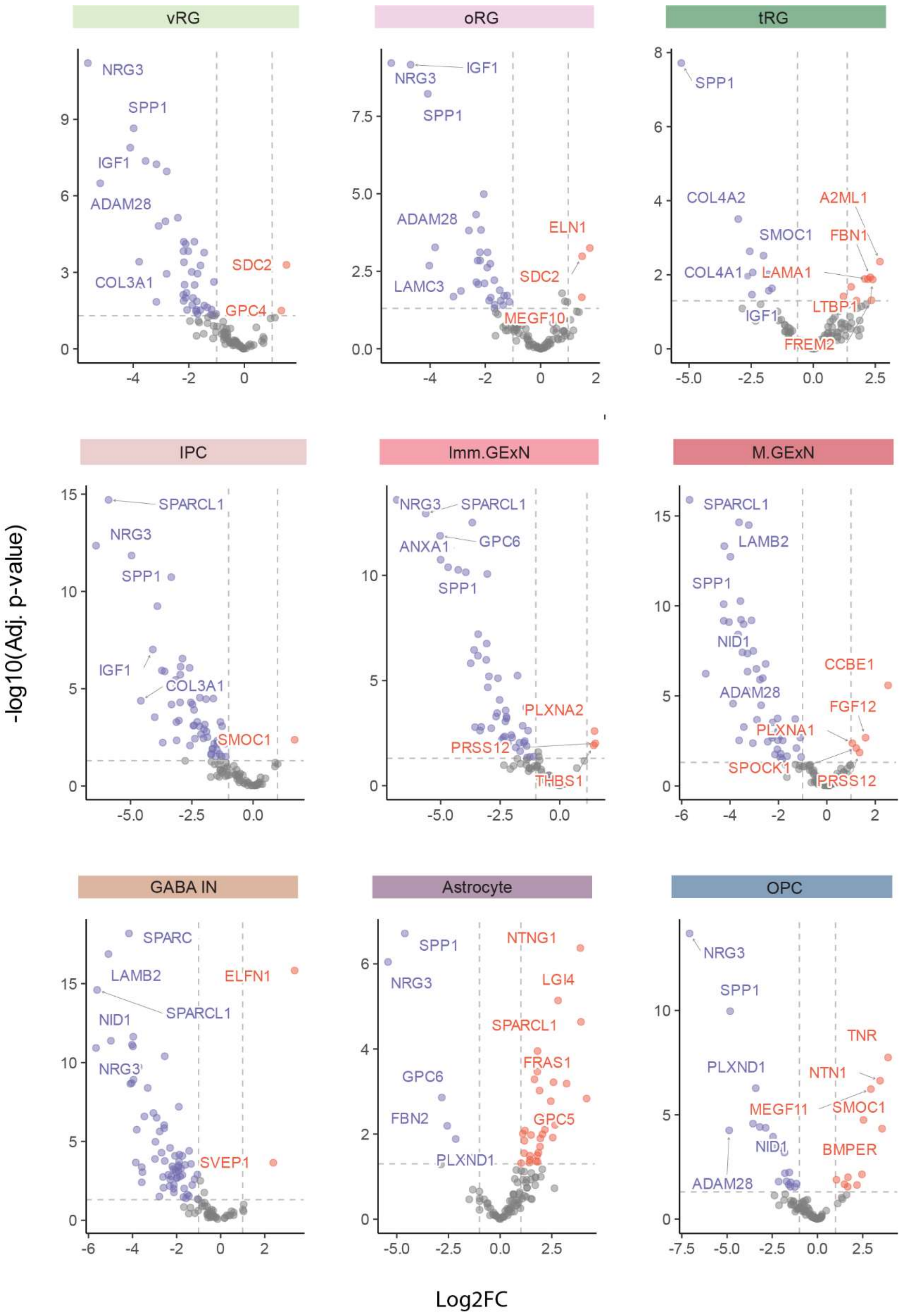


**Supplementary Fig. 6: Differentially expressed matrisome genes associated with NDDs**

Volcano plot illustrating differentially expressed matrisome genes in each cell type. The top five significantly upregulated and downregulated NDD associated matrisome genes are labeled (ranked by $\log_2$ fold-change >1, adjusted p-value < 0.05).
